# SeaMoon: from protein language models to continuous structural heterogeneity

**DOI:** 10.1101/2024.09.23.614585

**Authors:** Valentin Lombard, Dan Timsit, Sergei Grudinin, Elodie Laine

## Abstract

How protein move and deform determines their interactions with the environment and is thus of utmost importance for cellular functioning. Following the revolution in single protein 3D structure prediction, researchers have focused on repurposing or developing deep learning models for sampling alternative protein conformations. In this work, we explored whether continuous compact representations of protein motions could be predicted directly from sequences, without exploiting 3D structures. SeaMoon leverages protein Language Model (pLM) embeddings as input to a lightweight convolutional neural network. We assessed SeaMoon against ∼ 1 000 collections of experimental conformations exhibiting diverse motions. It predicts at least one ground-truth motion with reasonable accuracy for 40% of the test proteins. SeaMoon captures motions inaccessible to normal mode analysis, an unsupervised physics-based method relying solely on 3D geometry, and generalises to proteins without detectable sequence similarity to the training set. SeaMoon is easily retrainable with novel or updated pLMs.

## Introduction

Proteins coordinate and regulate all biological processes by adapting their 3D shapes to their environment and cellular partners. Deciphering the complexities of how proteins move and deform in solution is thus of utmost importance for understanding the cellular machinery. Yet, despite spectacular advances in protein structure determination and prediction, comprehending protein conformational heterogeneity remains challenging^1–3^.

Many recent approaches have concentrated on repurposing the protein structure prediction neural network AlphaFold2^4^ to generate conformational diversity^5^. Guiding the predictions with state-annotated templates proved successful for modelling the multiple functional states of a couple of protein families^6,7^. In addition, massive sampling strategies have shown promising results for protein complexes^8 9,10^ with notable success in the blind CASP15-CAPRI assessment^11^. While they can be deployed seamlessly with parallelized implementations^12^, they remain highly resource-intensive.

Other strategies have explored promoting diversity by modulating and disentangling evolutionary signals^13^. The rationale is that amino acid co-variations in evolution reflect 3D structural constraints^14–20^. These evolutionary patterns can be extracted directly from alignments of evolutionary related sequences, or, as shown more recently, by modeling raw sequences at scale with protein language models^21–23^. Inputting shallow, masked, corrupted or sub-sampled alignments to AlphaFold2 allowed for modelling distinct conformations for a few protein families^24–27^. Nevertheless, contradictory findings have highlighted difficulties in rationalising the effectiveness of these modifications and interpreting them, particularly for metamorphic proteins^28–30^.

More classically, physics-based molecular dynamics (MD) is a method of choice to probe protein conformational landscapes^31^. Nonetheless, the time scales amenable to MD simulations on standard hardware remain much smaller than those spanned by slow molecular processes^32^. This limitation has stimulated the development of hybrid approaches combining MD with machine learning (ML) toward accelerating or enhancing sampling^33^. Deep neural networks can help to identify collective variables from MD simulations as part of importance-sampling strategies^32,34–37^. Or they may directly generate conformations according to a probability distribution learnt from MD trajectories or sets of experimental structures^38–41^. Diffusion-based architectures^38,42,43^ and the more general flow-matching framework^44^ provide highly efficient and flexible means to generate diverse conformations conditioned on cellular partners and ligands. Nevertheless, they are prone to hallucination, and models trained across protein families still fail to approximate solution ensembles^42^.

On the other hand, the normal mode analysis (NMA) represents a data- and compute-inexpensive unsupervised alternative for accessing large-scale, shape-changing protein motions^45^. In particular, the NOLB method predicts protein functional transitions in real-time by deforming single structures along a few collective coordinates inferred with the NMA^46,47^. The generated conformations are physically plausible, stereochemically realistic, and some of them approximate known biologically relevant intermediate states^46^. However, the results strongly depend on the 3D geometry of the starting structure, and although some of the initial topological constraints can be easily alleviated^48^, the NMA remains unsuitable for modelling extensive secondary structure rearrangements.

Training and benchmarking predictive methods is difficult due to the sparsity and inhomogeneity of the available experimental data^49^. X-ray crystallography, cryogenic-electron microscopy (cryo-EM), and nuclear magnetic resonance spectroscopy (NMR) have provided invaluable insights into protein diverse conformational states^2,50^, but only for a relatively small number of proteins^51^. Small-angle X-ray or neutron scattering (SAXS, SANS) and high-speed atomic force microscopy (HS-AFM) techniques allow for directly probing continuous protein heterogeneity, but with limited structural resolution^52–54^.

Ongoing community-wide efforts aim at revealing the full potential of the available structural data by collecting, clustering, curating, visualising and functionally annotating experimental protein structures together with high-quality predicted models^50,55–61^. For instance, the DANCE method produces movie-like visual narratives and compact continuous representations of protein conformational diversity, interpreted as *linear motions*, from static 3D snapshots^62^. DANCE application to the Protein Data Bank (PDB)^49^ revealed that the conformations observed for most protein families lie on a low-dimensional *manifold*. Interpolation trajectories along the manifold can recapitulate known intermediate conformations, supporting its biological significance. Moreover, classical dimensionality reduction techniques can learn this manifold and generate unseen conformations with reasonable accuracy, albeit only in close vicinity of the training set ^62^.

Here, we explored the possibility of predicting protein motions directly from amino acid sequences without exploiting nor sampling protein 3D structures. To do so, we leveraged protein Language Models (pLMs) pre-trained through self-supervision over large databases of protein-related data. Our approach, SEAquencetoMOtioON or SeaMoon, is a 1D convolutional neural network inputting a protein sequence pLM embedding and outputting a set of 3D displacement vectors (**Fig. 1**). The latter define protein residues’ relative motion amplitudes and directions. We tested whether SeaMoon could capture the *linear motion manifold* underlying experimentally resolved conformations across thousands of diverse protein families^62^. To this end, we devised an objective function invariant to global translations, rotations, and dilatations in 3D space. SeaMoon achieved a predictive performance similar to the normal mode analysis (NMA) when inputting purely sequence-based pLM embeddings^23^ without any knowledge about protein 3D structures. It could generalise to proteins without any detectable sequence similarity to the training set and capture motions not directly accessible from protein 3D geometry. Injecting implicit structural knowledge with sequence-structure bilingual or multimodal pLMs^63,64^ further boosted the performance. This work establishes a community baseline and paves the way for developing evolutionary- and physics-informed neural networks to predict continuous protein motions.

**Figure 1:**
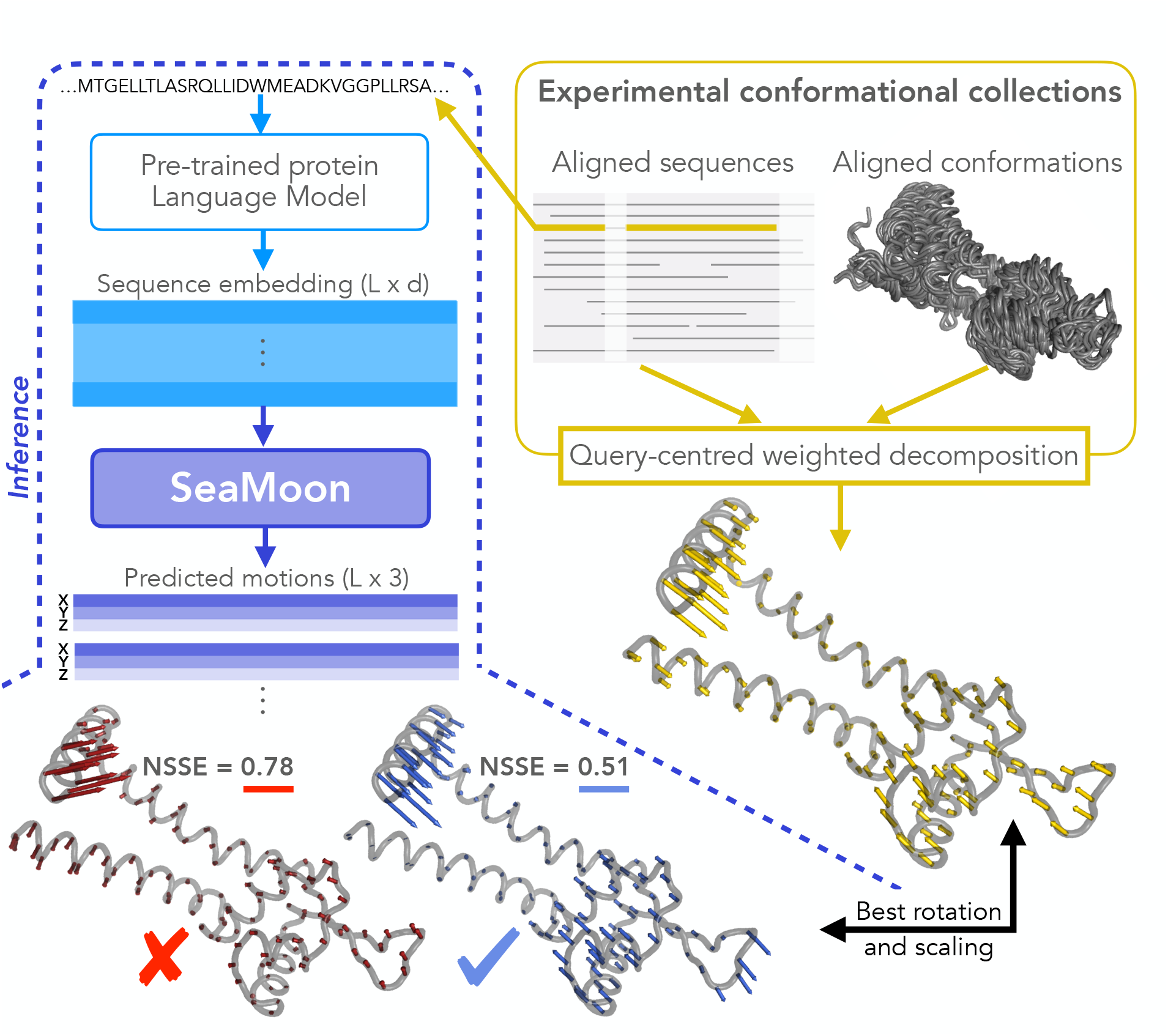
Outline of SeaMoon’s approach. SeaMoon takes as input a high-dimensional *L* × *d* matrix representation of a protein sequence of length *L* computed by a pre-trained pLM. It outputs a set of 3D vectors of length *L* representing linear motions. The training procedure regresses these output motions (blue and red arrows) against ground-truth ones (yellow arrows) extracted from experimental conformational collections through principal component analysis. For this, SeaMoon identifies the transformation (rotation and scaling) minimising their discrepancy, computed as a sum-of-squares error (SSE). We consider predictions with a normalised error (NSSE) smaller than 0.6 as acceptable. We show the query protein 3D structure only for illustrating the motions, it is not used by SeaMoon nor by the pLM generating the input embeddings.

## Results

### SeaMoon predicts protein motions from amino acid sequences alone

The approach introduced in this work, SeaMoon, predicts continuous representations of protein motions with a convolutional neural network inputting pLM sequence embeddings (**Fig. 1**). We trained and tested SeaMoon on over ∼ 17 000 experimental conformational collections representing a non-redundant set of the PDB at 80% sequence similarity. We used the principal components extracted from these collections as ground-truth linear motions to which we compared SeaMoon predicted 3D vectors three per protein by default). The latter are not anchored on a particular conformation and may be in any arbitrary orientation. To allow for a fair comparison, we determined the optimal rotation and scaling between the ground-truth and predicted vectors before computing a normalised sum-of-squares error (NSSE) between them (see *Methods* for details).

SeaMoon predicted protein motions with similar or better accuracy compared to the purely geometry-based unsupervised NMA when assessed against a test set of 1 121 proteins (**Fig. 2A** and **S1**). SeaMoon performance depends on the pLM used to compute the input embeddings from amino acid sequences. More specifically, the two structure-aware pLMs ESM3^63^ and ProstT5^64^ yielded a substantial improvement over the purely sequence-based pLM ESM2^23^. Furthermore, ProstT5 outperformed ESM3 (**Fig. 2A**, paired Wilcoxon signed-rank test p-values *<* 10^*−*6^), despite having a much smaller number of parameters and embedding dimensions (**Table S1**). ProstT5 is a fine-tuned version of the sequence-only model ProtT5 that translates amino acid sequences into sequences of discrete structural states and reciprocally, while ESM3 is a multi-modal pLM capable of conditioning on and reconstructing several protein sequence and structural properties. Beside the influence of the pLM, we observed a boost in performance by up to 10% upon stimulating the model to learn a one-sequence-to-many-motions mapping (**Fig. 2A**, compare plain and dotted lines). More specifically, we augmented the training data by using multiple (up to 5) reference conformations per experimental collection (**Table S2**). While the pLM embeddings within a collection should be highly similar, the extracted motions may differ substantially from one reference to another^62^. The positive impact of this data augmentation strategy was most visible for the ESM2-based version of SeaMoon (**Fig. 2A**).

**Figure 2:**
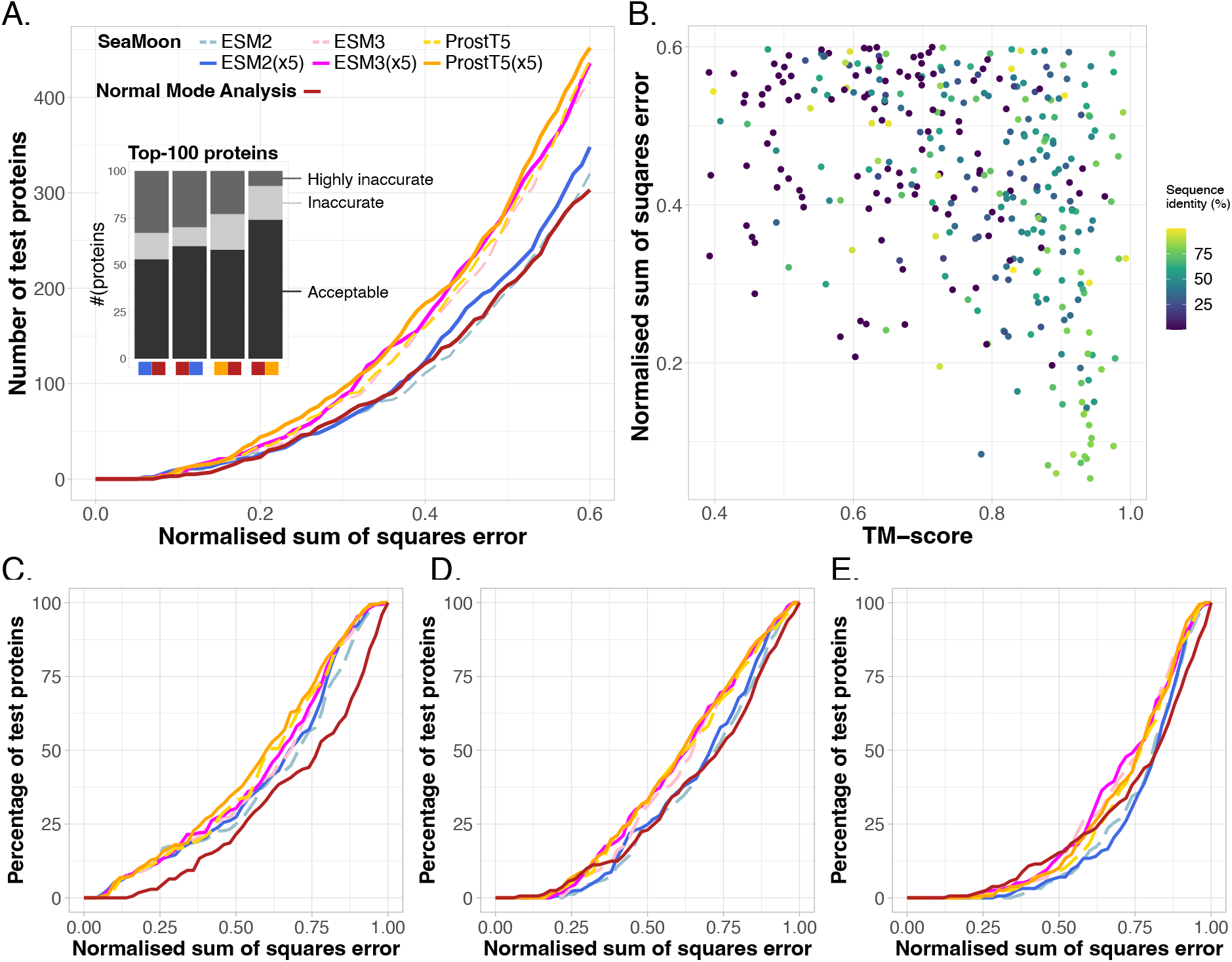
SeaMoon performance and generalisation capability. We report the NSSE of the best match between 3 predictions and 3 ground-truth motions for each test protein. **A**. Cumulative NSSE for six different versions of SeaMoon and for the NMA. We tested three pLMs, namely ESM2, ESM3 and ProstT5, and a data augmentation strategy with 5 training samples per experimental collection (x5). We cropped the plot at NSSE=0.6 for ease of visualisation; see Fig. S1 for the full curves. Inset: Agreement between a selection of methods. For instance, the first bar stack gives the numbers of proteins for which the NMA (right red square) produced acceptable (NSSE*<*0.6), inaccurate (0.6*<*NSSE*<*0.75) or highly inaccurate (NSSE*>*0.75) predictions among the top-100 proteins best-predicted by SeaMoon-ESM2(x5) (left blue square). **B**. NSSE computed for SeaMoon-ESM2(x5) in function of sequence and structural similarity to the training set; see Fig. S5 for the other SeaMoon versions and NMA. **C-E**. Cumulative NSSE computed on three subsets of test proteins with increasing difficulty. **A. Easy:** at least 70% sequence identity with at least one train protein. **B. Intermediate:** at most 30% sequence identity with any train set example and a TM-score of at least 0.7 with some train protein. **C. Difficult:** at most 30% sequence identity and 0.5 as TM-score with any train protein. The colour code is the same as in panel A.

Looking at individual predictions, one can appreciate how they degrade as the error (NSSE) increases (**Fig. 3**). A prediction with a NSSE smaller than 0.2 almost perfectly superimposes to the ground-truth motion (**Fig. 3**, 8E7MH). SeaMoon-ProstT5(x5) generated at least one such near-perfect prediction, among the three predicted, for 44 proteins, representing 4% of the test set (**Fig. 2A**). It achieved NSSE smaller than 0.4 for 184 proteins (16%) and smaller than 0.6 for 452 proteins (40%, **Table I**). We consider predictions with NSSE larger than 0.6 as inaccurate as they typically miss or indicate completely wrong directions for a large part of the residues involved in the motion (**Fig. 3**, see 4ZEVB, 6W19p, and 7RTNB). By comparison, the errors computed for random predictions are typically above 0.9 (**Fig. S1B**). Based on a quality NSSE cutoff of 0.6, SeaMoon reached the highest success rate of 40% with ProstT5, followed by ESM3 (39%) and ESM2 (31%), while the NMA success rate is 27% (**Table I**). Average success rates computed over 1 000 bootstrap subsampling simulations, each one containing ∼ 100 test proteins, deviate by less than 0.5% from the estimates computed over the full test set (**Fig. S2**). The corresponding distributions follow the same trend as the cumulative error curves (**Fig. 2A** and **S1**). In particular, the interquartile range at a NSSE cutoff of 0.6 is 37-44% for SeaMoon-ProstT5(x5), significantly higher than 24-30% for the NMA (**Fig. S2**).

**Table I:** Performance and dependence on the similarity to the training set.

| Method | Protocol | Number of proteins with acceptable predictions (NSSE<0.6) |  |  |  |
| --- | --- | --- | --- | --- | --- |
|  |  | overall | easy | intermediate | difficult |
| Total |  | 1 121 | 172 | 161 | 158 |
| SeaMoon | ESM2 | 320 (29%) | 66 (38%) | 53 (33%) | 25 (16%) |
|  | ESM2(x5) | 348 (31%) | 68 (40%) | 58 (36%) | 21 (13%) |
|  | ESM3 | 416 (37%) | 70 (41%) | 66 (41%) | **43 (27%)** |
|  | ESM3(x5) | 436 (39%) | 73 (42%) | 75 (47%) | 41 (26%) |
|  | ProstT5 | 439 (39%) | 84 (49%) | 76 (47%) | 29 (18%) |
|  | ProstT5(x5) | **452 (40%)** | **86 (50%)** | **77 (48%)** | 37 (23%) |
| NMA |  | 303 (27%) | 57 (33%) | 57 (35%) | 35 (22%) |

**Figure 3:**
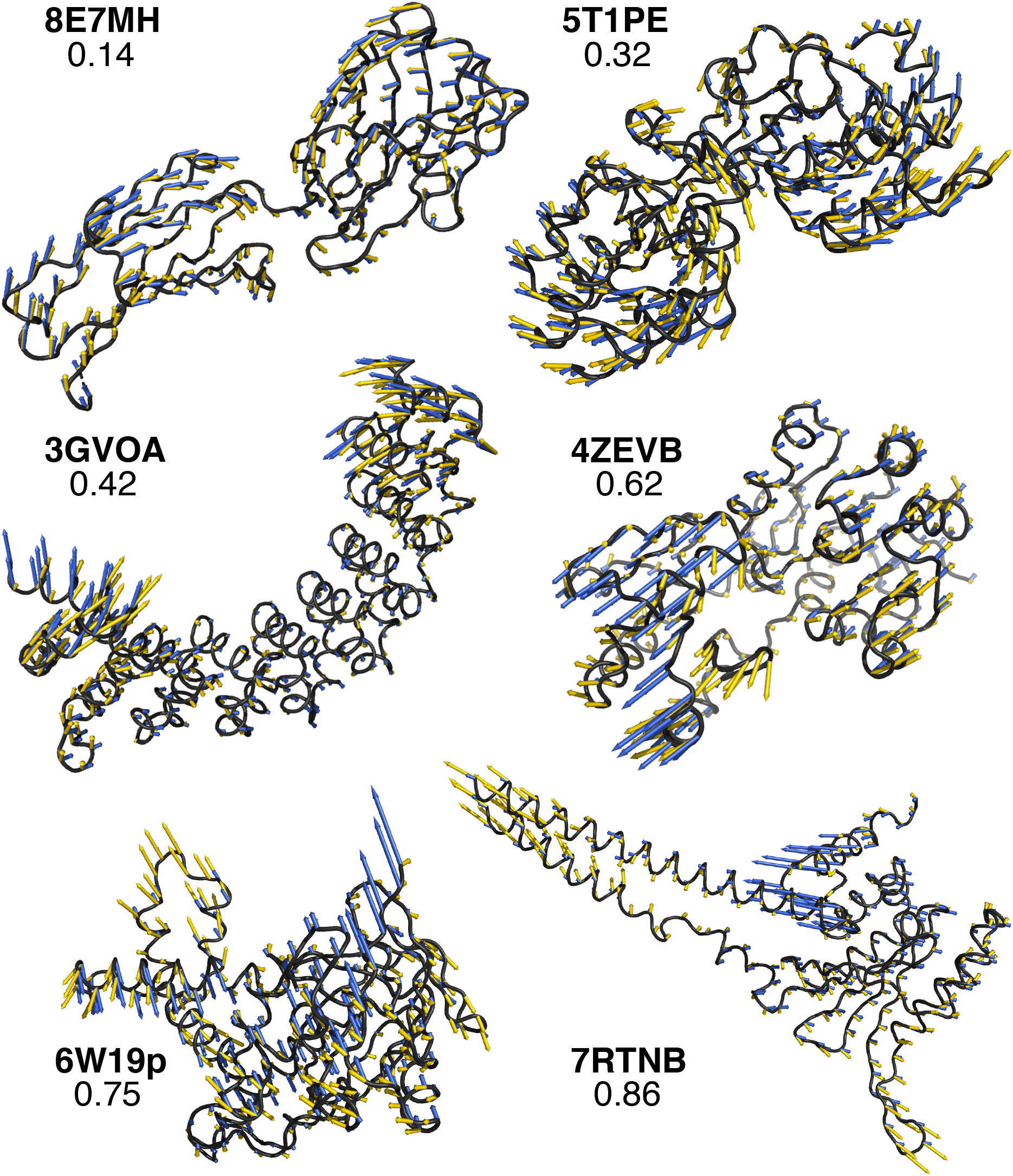
Examples of predictions. They allow for a visual assessment of how well the predicted vectors (in blue) approximate the ground-truth motions (in yellow) at different levels of NSSE (indicated on each panel). For each example, the query conformation is shown in black cartoons and labelled with its PDB chain identifier (in bold). We obtained the predicted vectors with SeaMoon-ESM2(x5).

### SeaMoon generalises to unseen proteins across diverse protein families

SeaMoon produced high-quality predictions at different levels of similarity to the training set, which we can interpret as varying difficulty levels (**Table I, Fig. 2B**, and **Fig. S3**). It generated acceptable predictions for up to 50% of the easy cases, namely the test proteins sharing over 70% sequence similarity with at least one train protein (**Table I**). The predictions are almost perfect (NSSE *<* 0.2) for about 20 proteins, representing 10% of this subset (**Fig. 2C**). Most of them are antibodies, a class of proteins well represented in both train and test sets (**Fig. 4A**). Beyond such easy cases, SeaMoon achieved similar success rates on test proteins sharing similar 3D folds with some train proteins (TM-score*>*0.7), despite highly divergent sequences, below 30% identity (**Table**

**Figure 4:**
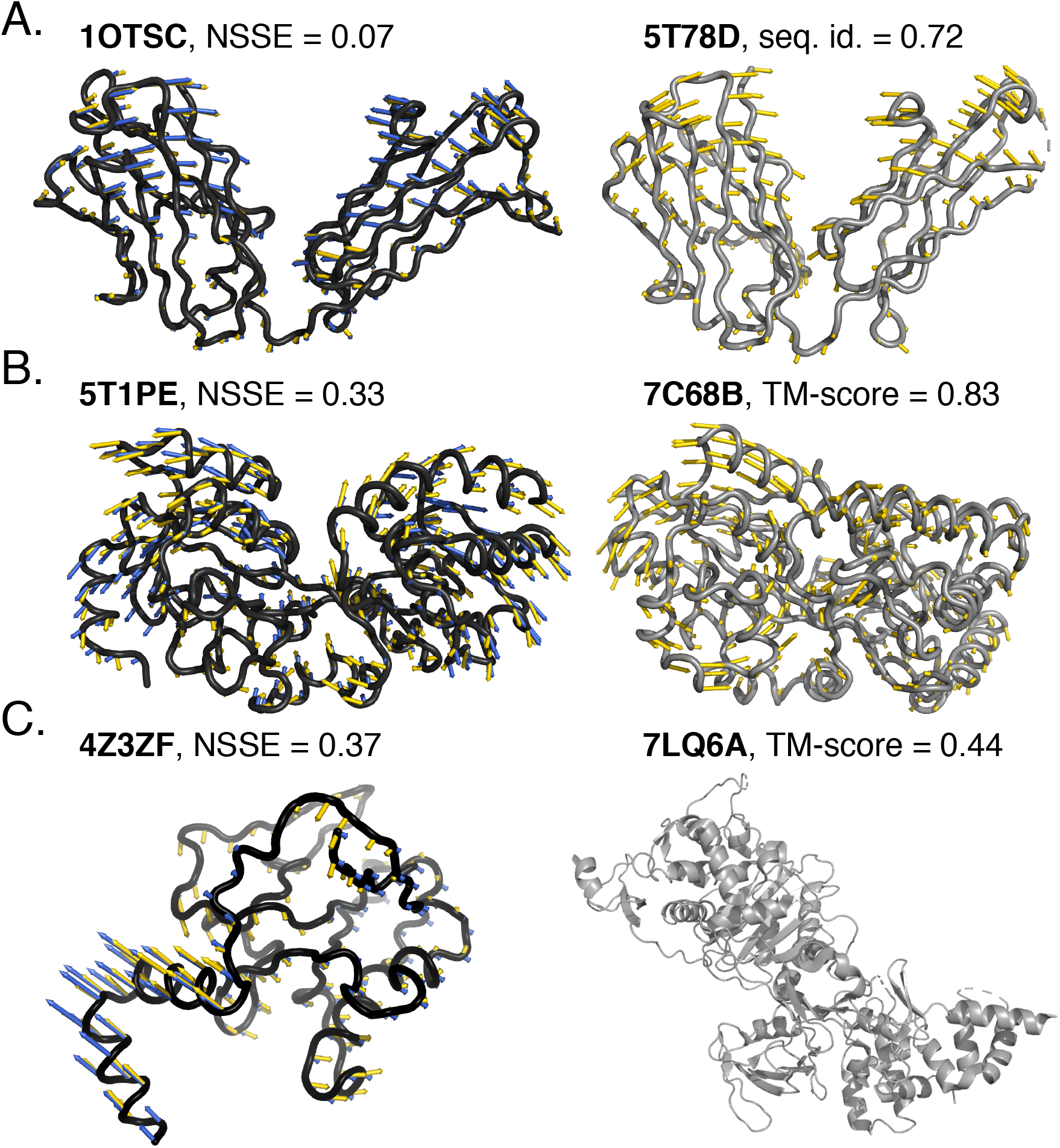
Examples of predictions for test proteins with decreasing similarity to the training set. The conformations are shown in cartoons and labelled with their PDB chain identifier. The ground-truth and SeaMoon-ESM2(x5) predicted motions are depicted with yellow and blue arrows, respectively. Left, in black: test proteins. Right, in grey: closest proteins from the training set. **A**. Fab fragment (heavy chain), 221 residues, 107 conformations in the collection, collectivity *κ* = 0.74 for the ground-truth motion. Its sequence, structure and main motion are highly similar to the Fab fragment displayed on the right **B**. Putative ABC transporter from *Campylobacter jejuni*, 326 residues, 8 conformations, *kappa* = 0.74. It does not have any detectable sequence similarity to the training set. Its structure and main motion bear some resemblance with the ABC transporter from *Thermus thermophilus* shown on the right. **C**. Iron-sulfur cluster-binding oxidoreductase, 170 residues, 20 conformations, *kappa* = 0.52. It does not have any detectable sequence similarity to the training set and the structurally closest training protein, the bacterial penicillin-binding protein 1B, exhibits a different 3D fold and different motions.

**I** and **Fig. 2D**). These test proteins represent an intermediate level of difficulty for which the ATP-binding cassette (ABC) transporter superfamily provides an illustrative example. In particular, SeaMoon-ESM2(x5) successfully transferred the opening-closing motion characteristic of the “Venus Fly-trap” mechanism for transporting sugars^65^ from ABC transporters in the training set to the held-out putative ABC transporter from *Campylobacter jejuni* (**Fig. 4B**, 5T1PE, NSSE=0.33). The latter does not have any detectable sequence similarity with any train protein but shares high structural similarity with the sugar ABC transporter from *Thermus thermophilus* (7C68B, TM-score = 0.83). Finally, while SeaMoon predictions tend to display higher errors on the most challenging subset, defined with TM-score below 0.5 and sequence similarity below 30%, its success rate at NSSE=0.6 remains comparable or better than that of the NMA (**Table I** and **Fig. 2E**). Successful cases completely unrelated to the training set include the benzoyl-coenzyme A reductase from *Geobacter metallireducens* (**Fig. 4C**, 4Z3ZF). SeaMoon-ESM2(x5) achieved NSSE=0.37 on this protein, whose structurally closest train protein, the bacterial penicillin-binding protein 1B (7LQ6A, TM-score=0.44), exhibits a different 3D fold and different motions.

The TM-align algorithm^66^, which we used to compute TM-scores, compares protein structures by treating them as rigid bodies that cannot deform. This approach might underestimate the similarity between structures that actually represent the same fold in different conformational states. To address this limitation, we re-evaluated the TM-scores between the most structurally similar train-test protein pairs using Kpax flexible fragment-based structural alignment algorithm^67^. Unlike TM-align, Kpax can accommodate conformational changes by aligning protein fragments independently. The cumulative NSSE loss curves obtained on the Kpax-based intermediate and difficult subsets show the same trends as those computed from the TM-align-based subsets (**Fig. S4**). This analysis confirms that SeaMoon is able to recognise previously seen folds and generalise to unseen ones. Furthermore, while the structure-aware pLMs consistently yield the best performance across the different categories, ESM3 becomes more advantageous compared to ProstT5 as the similarity to the training set decreases (**Table I, Fig. 2**, and **Fig. S4**). This observation suggests that ESM3 larger parameter count and access to functional properties during pre-training may provide it with better generalisation capabilities.

### SeaMoon is complementary to the normal mode analysis

We observed a substantial overlap between the sets of successful predictions generated by SeaMoon and NMA, as well as between their respective NSSE distributions (**Fig. 2A**, inset, and **Fig. S5**). Specifically, SeaMoon base version with ESM2 embeddings generated acceptable predictions for 60% of the top-100 test proteins best-predicted by the NMA, and this proportion reaches 75% with ProstT5 embeddings (**Fig. 2A**, inset). Using implicit structural knowledge allowed recovering elastic motions, such as that exhibited by the mammalian plexin A4 ectodomain (**Fig. S6**, 5L5LB, NSSE=0.28). Reciprocally, about half of the top-100 proteins best-predicted by SeaMoon exhibit motions accessible to the NMA (**Fig. 2A**, inset).

Most of the motions well captured by the two approaches involve a large portion of the protein and correspond to large conformational changes. They include functional opening-closing motions of virulence factors, thermophilic proteins, metalloenzymes, periplasmic binding proteins, dehydrogenases, glutamate receptors, and antibodies (**Fig. S7**). To further validate SeaMoon’s ability to predict a wide range of opening-closing motions, we evaluated its performance on the iMod benchmark^68^, which was specifically designed to assess coarse-grained elastic network models (**Fig. S8**). SeaMoon-ProstT5(x5) matched the success rate of the NMA on this dataset, capturing the motions of 12 out of 21 proteins with high accuracy (NSSE*<*0.4), including adenylate kinase hinge motion (1LAFE) and the chaperone GroEL complex motion (1AONA). In addition, SeaMoon-ProstT5(x5) outperformed the NMA on the two shear motions in the dataset, exhibited by the *E. coli* DNA Polymerase III (1MMIA) and the adenosyl-cobinamide kinase (1C9KA), and on the allosteric motion of Aspartate carbamoyltransferase (1EKXA).

Despite the overall agreement between the two approaches, the NMA performed extremely poorly for a quarter to a third of SeaMoon top-100 test proteins (**Fig. 2A**, inset). The associated motions tend to be localised, with a median value of collectivity *κ* = 0.20. More broadly, we confirmed this trend by statistically assessing whether the set of motions well captured by the NMA were enriched or depleted in motions of different sizes (**Fig. 5**). The set of acceptable NMA predictions contains about twice more global motions (*κ >* 0.60) than the full test set and four times fewer localised motions (*κ <* 0.30, **Fig. 5A**). By contrast, the sets of acceptable SeaMoon predictions do contain roughly the same proportions of localised motions as in the full test set. These results indicate that SeaMoon is more capable of handling localised motions than the NMA. The bacterial toxin PemK provides an illustrative example of this capability (**Fig. 6A**). SeaMoon-ESM2(x5) captured the PemK’s loop L12 motion with high precision (**Fig. 6A**, NSSE=0.24) whereas the NMA failed to delineate the mobile region in the protein and to infer its direction of movement (**Fig. 6A**, in red). This highly localised motion (*κ* = 0.17) plays a decisive role in regulating PemK RNAse activity by promoting the formation of the PemK-PemI toxin-antitoxin^69^.

**Figure 5:**
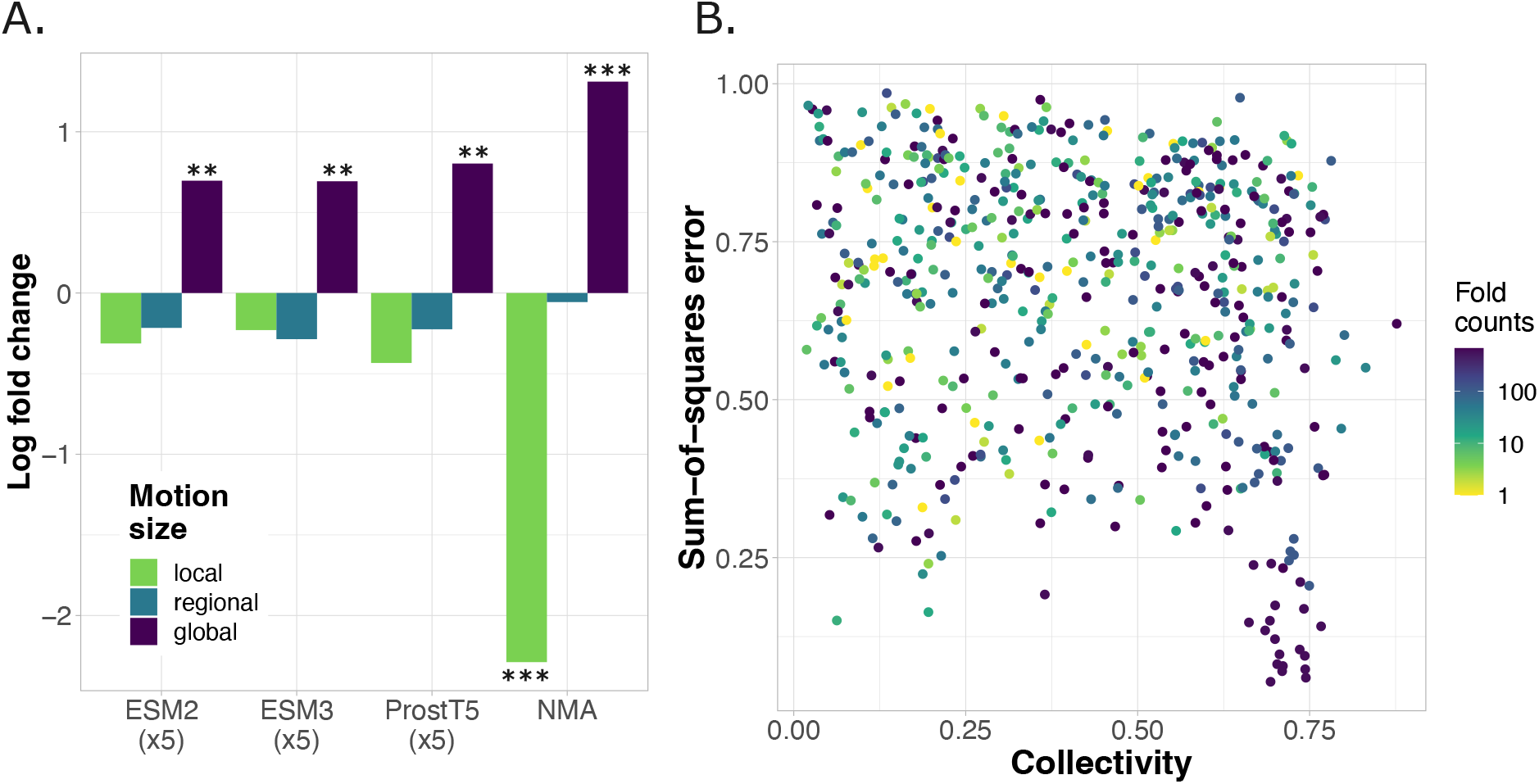
SeaMoon performance depending on the type of motion and fold representativity. A. Log_2_ fold change in the proportion of local, regional and global motions between the set of acceptable predictions (NSSE*<*0.6), for each method, and the full test set. The categories of motions are defined based on collectivity computed on the ground-truth motions. Local: *κ*≤ 0.30. Regional: 0.30 *< κ* ≤ 0.60. Global: *κ >* 0.60. The stars indicate the significance of the enrichment or depletion, as estimated by a hypergeometric test, namely ** for a p-value below 10^*−*8^ and *** below 10^*−*40^ (see also **Table S3**). **B**. NSSE computed for SeaMoon-ESM2(x5) in function of motion collectivity and fold (CATH topology) counts in the training set. Each dot correspond to a protein and we report collectivity for the ground-truth motion best approximated by the predictions. Only test proteins with at least one annotated CATH domain are shown. In case of multiple domains, we consider the maximum fold count value.

**Figure 6:**
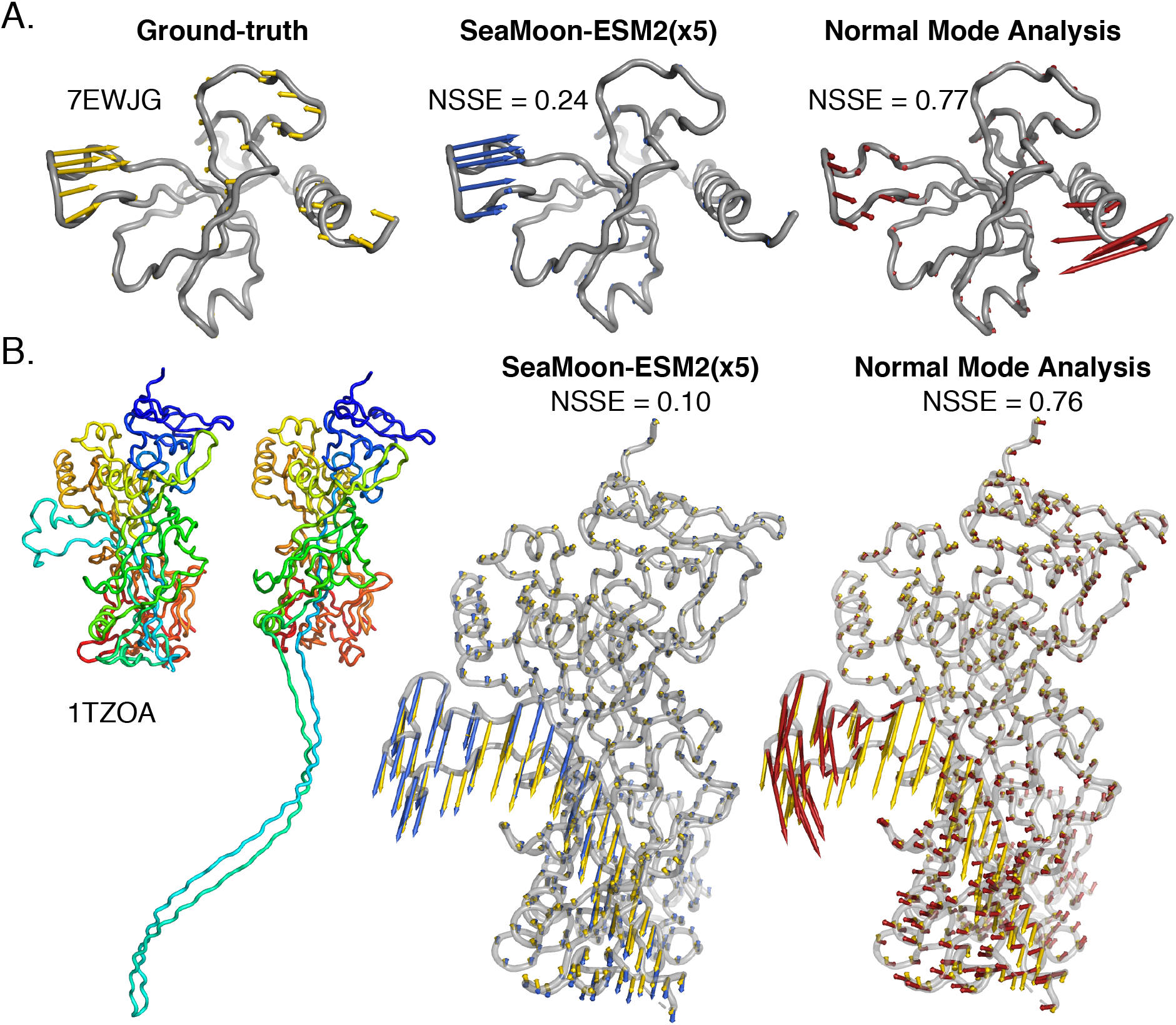
Examples of motions well predicted by SeaMoon and not by the NMA. The arrows depicted in yellow, blue and red on the grey 3D structures represent the ground-truth motions and the best-matching predictions from SeaMoon-ESM2(x5) and the NMA, respectively. **A**. Bacterial toxin PemK (PDB code: 7EWJ, chain G) from the test set. It does not have any detectable sequence similarity to the training set **B**. Anthrax protective antigen (PDB code: 1TZO, chain A) from the validation set. We show the two most extreme conformations of the collection on the left, colored according to the residue index, from the N-terminus in blue, to the C-terminus in red. The closest homolog from the training set shares 35% sequence similarity.

In addition, SeaMoon allows to go beyond the elastic approximation of the NMA and hence, may be a better option for modeling deformations that imply large secondary structure rearrangements, such as fold-switches. This advantage is revealed by the protective antigen (PA) from anthrax (**Fig. 6B**). SeaMoon-ESM2(x5) accurately predicted the relative motion amplitudes and directions of an 80 residue-long region that detaches from the rest of the protein upon forming an heptameric pore **Fig. 6B**). By contrast, the NMA predicted a breathing motion poorly approximating the ground-truth one (**Fig. 6B**). PA’s ∼ 30°A-large conformational transition is essential for the translocation of the bacterium’s edema and lethal factors to the host cell^70^. Neither PemK nor PA have any detectable sequence similarity to the training set. SeaMoon likely leveraged information coming from training proteins with similar folds and functions from other bacteria^71,72^.

### SeaMoon can recapitulate entire motion subspaces

Beyond assessing individual predictions, we evaluated the global similarities between predicted and ground-truth 3-motion subspaces focusing on the test proteins for which SeaMoon produced at least one acceptable prediction (**Table I**). We found that SeaMoon motion subspaces were fairly similar to the ground-truth ones, with a Root Mean Square Inner Product (RMSIP)^73–75^ higher than 0.5, for almost two thirds of these proteins. We observed an excellent correspondence for a dozen proteins, *e*.*g*., the *Mycobacterium* phage Ogopogo major capsid protein (**Fig. 7** and **Fig. S9**). The purely sequence-based SeaMoon-ESM2(x5) achieved an RMSIP of 0.75 on this protein, and the structure-aware SeaMoon-ProstT5(x5) reached 0.82. SeaMoon-ProstT5(x5) first, second and third predicted motions had a Pearson correlation of 0.93, 0.73 and 0.75 with the first, third and second ground-truth principal components, respectively (**Fig. 7A**). The associated NSSE were all smaller than 0.5 (**Fig. 7B**). By inspecting the training set, we could identify several major capsid proteins from other bacteriophages sharing the same HK97-like fold as the Ogopogo one (TM-score up to 0.78), despite relatively low sequence similarity (up to 34%). The ability of SeaMoon to recapitulate the Ogopogo protein entire motion subspace with reasonable accuracy likely reflects the high conservation of major capsid protein dynamics upon forming icosahedral shells^76^.

**Figure 7:**
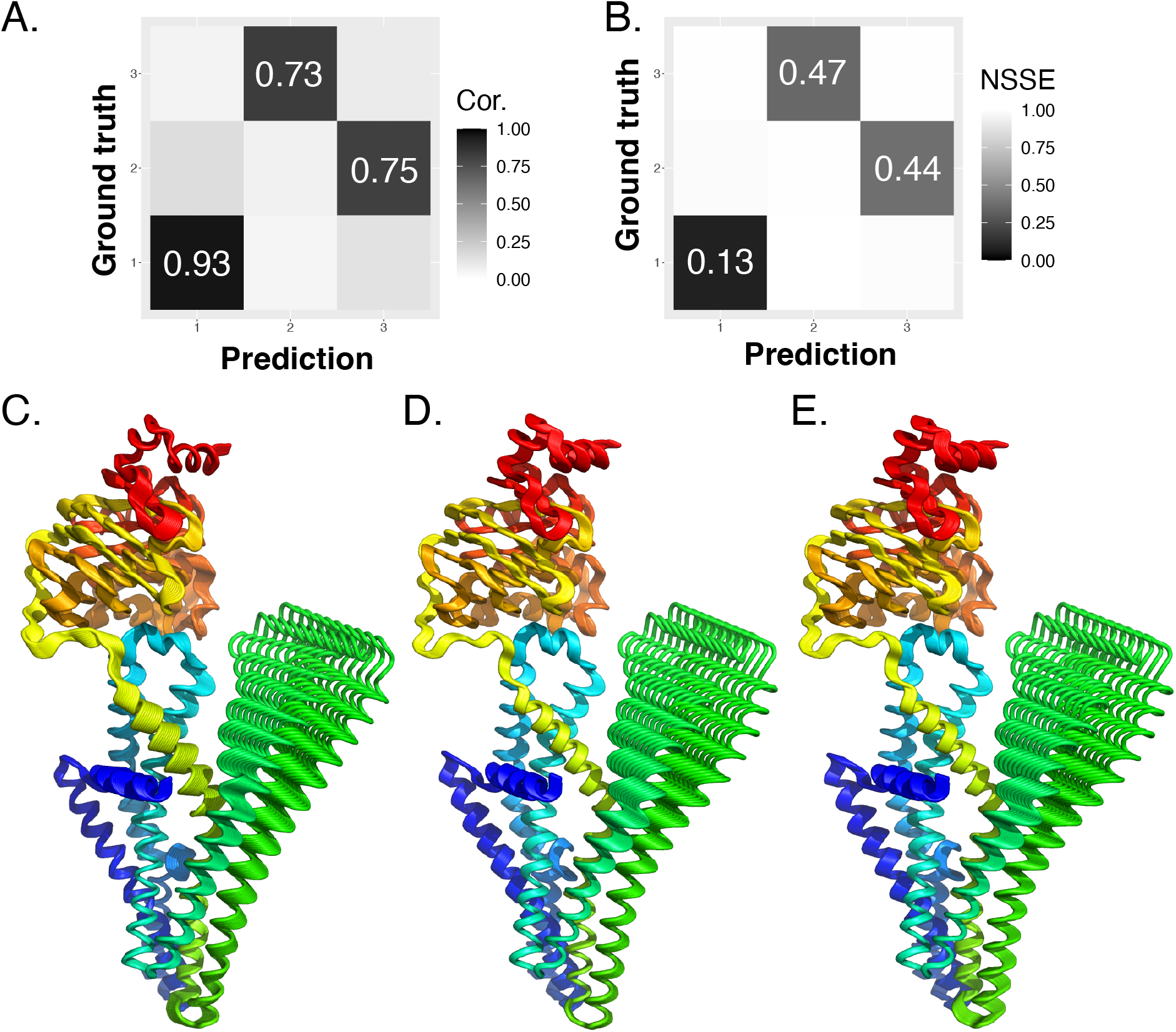
Motion subspace comparison and deformation trajectories. A-B. Ogopogo major capsid protein motion subspace. PDB code: 8ECN, chain B. **A**. Pairwise similarities measured as Pearson correlations between the ground-truth motions and SeaMoon-ProstT5(x5) predictions. **B**. Pairwise discrepancies measured as NSSE. **C-E**. Trajectories of a human ABC transporter (PDB code: 7D7R, chain A) deformed along its first ground-truth principal component (C) and the best-matching SeaMoon-ProstT5(x5) prediction (D-E). **D**. The prediction is optimally aligned with the ground truth. **E**. The orientation of the prediction minimises the protein conformation’s angular velocity. Each trajectory comprises 10 conformations coloured from blue at the N-terminus to red at the C-terminus.

### Influence of fold representativity

Protein folds are not evenly represented in the PDB, with some folds being significantly more abundant than others. Yet, we chose not to correct for this bias in our PDB-wide training set of conformational collections, because the same fold may exhibit different motions in different collections. This design choice raises two important questions. First, does SeaMoon performance remain consistent on a redundancy-reduced version of the test set? To address this, we computed average best-matching motion pair errors for each of the 252 folds represented in the test set, using CATH topologies as fold definitions^77^. This analysis showed that the per-fold performance estimates align well with those computed at the protein level (**Fig. 8**, compare panels A and B). The success rate computed for SeaMoon-ProstT5(x5) is 37% (at NSSE*<*0.6) compared to 40% without redundancy reduction. Moreover, SeaMoon performance remains similar to or better than the NMA (**Fig. 8A-B**). The second question concerns whether the motions of the folds that are over-represented in the training set are consistently better captured than those of under-represented folds (**Fig. 5B**). We tested this hypothesis by fitting a linear regression between the per-fold NSSEs and the fold frequency counts in our training set. We found no significant association between fold representativity and the per-fold average NSSEs, and only a weak association with the per-fold minimum NSSEs (**Fig. 8C-D**, adjusted R^2^ of 0.20 for SeaMoon-ProstT5(x5)). These results demonstrate that SeaMoon predictions are not significantly biased by the unbalanced representation of protein folds in the training set. Furthermore, SeaMoon-ProstT5(x5) success rate at NSSE*<* 0.6 computed on the test proteins that do not share any fold with the training set is 43% (12 out of 28 proteins), similar to the success rate computed over the full test set (40%).

**Figure 8:**
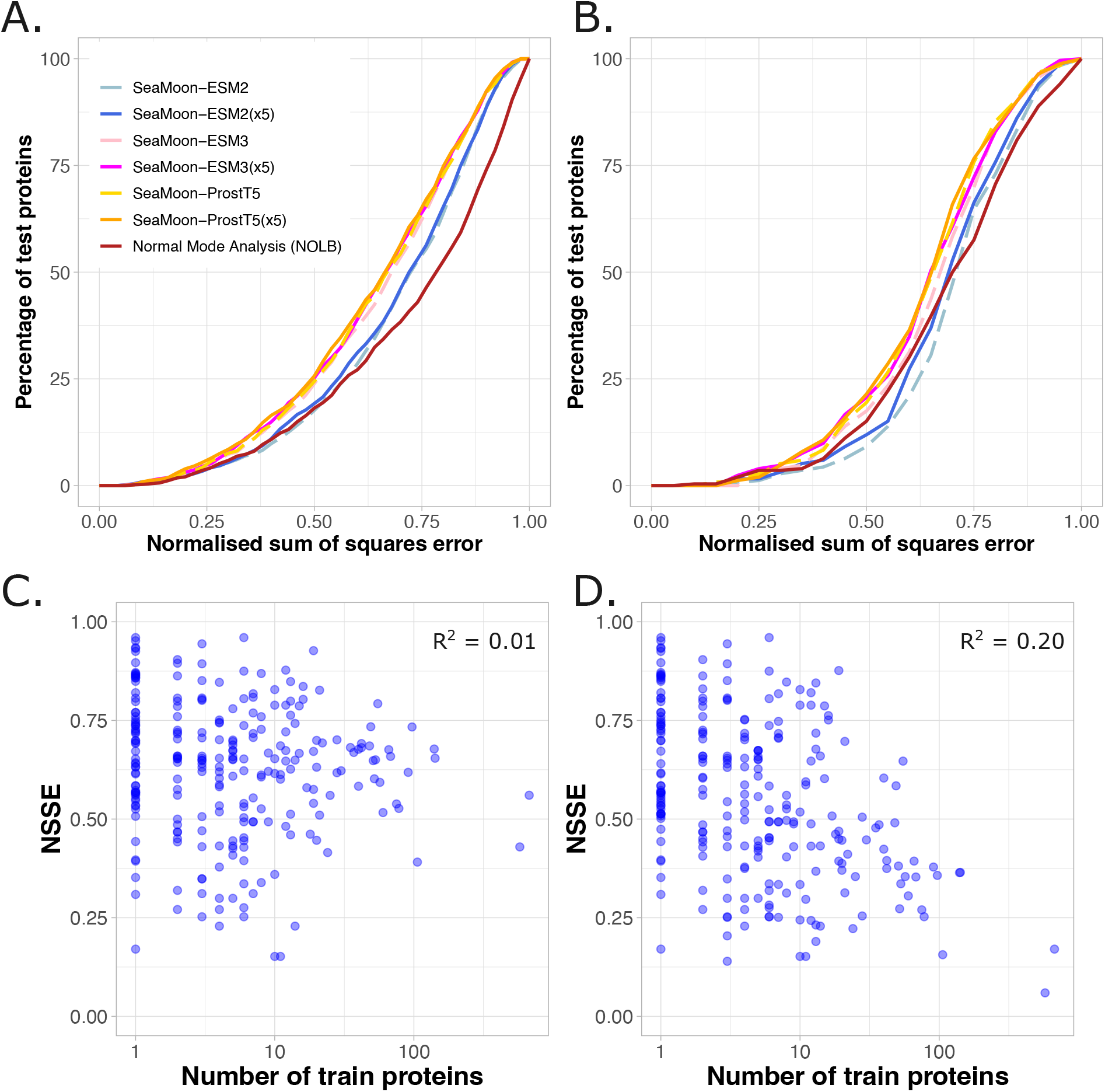
Influence of fold representativity. A-B. Cumulative NSSE computed per protein (A) and per fold (B), defined as topology in the CATH classification. The per-fold NSSEs are computed as the average NSSEs over the proteins associated to each fold. **C-D**. Per-fold NSSE in function of the number of train proteins containing at least one domain from this fold. The per-fold NSSEs are computed as the average NSSEs (C) of the minimum NSSEs (D) over the proteins associated to each fold. We report the adjusted R-squared values estimated by fitting a linear regression between the per-fold NSSEs and the log of the per-fold train protein counts.

### Contributions of the inputs and design choices

We investigated the contribution of SeaMoon inputs, architecture and objective function to its success rate through an ablation study, starting from SeaMoon-ProstT5 baseline model (**Table S4** and **Fig. 9**). Inputting random matrices instead of pre-trained pLM embeddings or using only positional encoding had the most drastic impacts. Still, we observed that the network can produce accurate predictions for over 100 proteins in this extreme situation (**Fig. 9**, in grey). Annihilating sequence embedding context by setting all convolutional filter sizes to 1 also had a dramatic impact, reducing to success rate from 40 to 25% (**Table S4** and **Fig. 9**). Moreover, a 7-layer transformer architecture (see *Methods*) underperformed SeaMoon’s convolutional neural network, despite having roughly the same number of free parameters (**Fig. 9**, in brown). Finally, disabling either sign flip or reflection (*i*.*e*., pseudo-rotation) or permutation when computing the loss degraded the performance by 6 to 15% (**Fig. 9**, in light green). This result underlines the utility of implementing a permissive and flexible comparison of the predicted and ground-truth motions during training.

**Figure 9:**
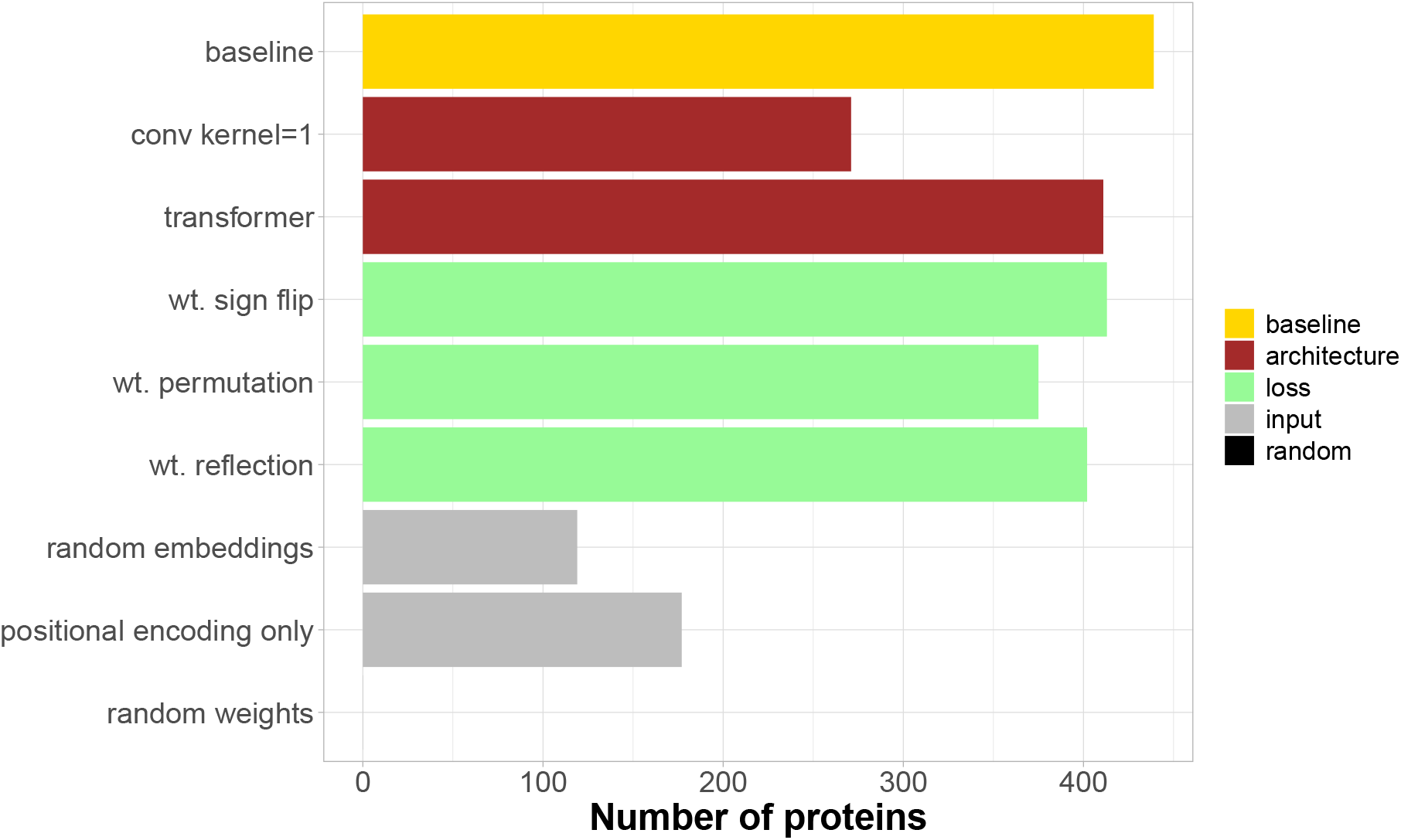
Ablation study. We report the number of test proteins with at least one acceptable prediction. The baseline model is SeaMoon-ProstT5.

### SeaMoon practical utility to deform protein structures

SeaMoon does not use any explicit 3D structural information during inference. Its predictions are independent of the global orientation of any protein conformation, making it impractical to directly use them to deform protein structures. To partially overcome this limitation, we propose an unsupervised procedure to orient SeaMoon predicted vectors with respect to a given protein 3D conformation. This method exploits the rotational constraints of the ground-truth principal components. Namely, the total angular velocity of the reference conformation subjected to a ground-truth principal component is zero (see *Methods*). Therefore, we determine the rotation that must be applied to the predicted motion vectors to minimize the total angular velocity of a target conformation.

This strategy proved successful for the vast majority of SeaMoon’s highly accurate predictions. SeaMoon-ProstT5(x5) predicted motion vectors, oriented to minimise angular velocity, exhibit an acceptable error (*<* 0.6) in 85% of cases where the optimal alignment with the ground truth results in NSSE *<* 0.3. This result indicates that predictions that approximate well the ground-truth principal components also preserve their properties. The human ABC transporter sub-family B member 6 gives an illustrative example where the third predicted motion vector approximates the first ground-truth principal component with NSSE = 0.20 upon optimal alignment and 0.22 upon angular velocity minimisation (**Fig. 7C-E**). Overall, the procedure allowed for correctly orienting acceptable predictions for 215 test proteins.

Note that this post-processing increases computing time significantly, from 12s to 24m over the 1 121 test proteins on a desk computer equipped with Intel Xeon W-2245 @ 3.90 GHz.

## Discussion

This proof-of-concept study explores the extent to which protein sequences encode functional motions. SeaMoon reconstructs these motions within an invariant subspace directly from pLM embeddings generated from amino acid sequences alone. Our results demonstrated two key findings. First, pLMs that had been exposed to structural information during their pre-training outperformed those trained exclusively for sequence reconstruction. Second, they highlighted SeaMoon’s ability to transfer knowledge about motions across distant homologs, leveraging the universal representation space of pLMs.

SeaMoon’s transfer learning approach makes it suitable for systematically assessing the evolutionary conservation of protein motions. Moreover, it complements unsupervised methods that rely entirely on the 3D geometry of protein structures, such as Normal Mode Analysis (NMA). Additionally, SeaMoon is highly computationally efficient, and thus applicable on a large scale. It took 12s to predict 3 motions for each of 1 121 test proteins on a desk computer equipped with Intel Xeon W-2245 @ 3.90 GHz.

One current limitation is the scarcity of functional motions in the training set, raising concerns about its accuracy and completeness. Both SeaMoon and NMA struggle to predict certain motions, suggesting that these may lack biological or physical relevance. They may result from experimental artifacts, most often of crystallographic origin. Nevertheless, establishing objective criteria to reliably distinguish artifactual conformations from biologically relevant ones remains infeasible. Our working assumption is that a subset of the conformational manifold in a protein collection always represents functional motions. To address this challenge, we designed our training loss function specifically to evaluate submanifolds by calculating the minimum error between each reference motion and the set of predicted motions, allowing the model to capture conformational diversity while mitigating the impact of potential artefacts.

Future work will explore incorporating explicit structural information. This should resolve the ambiguities in orienting predicted motions at inference time and could unlock SeaMoon’s potential for sampling or deforming conformations. Furthermore, we will test the potential of augmenting both training and evaluation sets with *in silico* generated data. This could include motions derived from MD and NMA simulations, as well as protein conformations predicted by AlphaFold. In addition, future improvements will include more explicit descriptions of the protein environment. For instance, we plan to condition predictions on the presence of other molecules. This is particularly important because only 15% of all 750K protein chains available in the PDB are monomeric^62^. Consequently, motions induced by binding partners or ligands likely constitute a significant portion of our training dataset.

Despite these limitations and avenues for improvement, the current findings offer valuable insights for integrative structural biology. SeaMoon provides compact representations of continuous structural heterogeneity in proteins. Such representations have high interpretability and explanatory power for most of the cases. The estimated motion subspaces can readily be used to compute protein conformational entropy. Lastly, the SeaMoon framework is highly versatile, featuring a lightweight, trainable deep learning architecture that does not depend on fine-tuning a large pre-trained model. This flexibility allows users to easily adapt the system to new input pLM embeddings without modifying the model architecture.

## Acknowledgments

The Sorbonne Center for Artificial Intelligence (SCAI) provided a salary to VL and computational resources. The authors thank Institut de Biologie Paris-Seine (IBPS) at Sorbonne Université for funding via a Collaborative Grant (Action Incitative) to DT. This work was co-funded by the European Union (ERC, PROMISE, 101087830). Views and opinions expressed are however those of the author(s) only and do not necessarily reflect those of the European Union or the European Research Council. Neither the European Union nor the granting authority can be held responsible for them. For the purpose of Open Access, a CC-BY public copyright licence has been applied by the authors to the present document and will be applied to all subsequent versions up to the Author Accepted Manuscript arising from this submission.

## Author Contributions

S.G. and E.L. designed research and supervised the project. V.L. designed the model’s architecture and carried out its implementation. S.G. and D.T. wrote the proofs and problem formalisation for orienting predictions with respect to a protein conformation with feedback from E.L.. D.T. implemented the solver. V.L., E.L. and S.G. produced and analysed the results. E.L. wrote the manuscript with input, support and feedback from all authors. All authors edited, read, and approved the final manuscript.

## Declaration of Interests

The authors declare no competing interests.

## Methods

### Data and code availability

The source code and model weights of this work are freely available at https://github.com/PhyloSofS-Team/seamoon. The data used for development and evaluation of SeaMoon are freely available at Zenodo^78^.

### Methods details

#### Datasets

To generate training data, we constructed a non-redundant set of conformational collections representing the whole PDB (as of June 2023) using DANCE^62^. To ensure high quality of the data, we replaced the raw PDB coordinates with their updated and optimised versions from PDB-REDO whenever possible^79^. We used a stringent setup where each conformational collection is specific to a set of close homologs. Specifically, any two protein chains belonging to the same collection share at least 80% sequence identity and coverage. We filtered out the collections with too few or too many data points. Namely, we asked for at least 5 conformations and a representative protein chain comprising between 30 and 1 000 residues. We further retained only C*α* atoms (option -c) and used coordinate weights to account for uncertainty (option -w).

For each collection, DANCE extracted the *K* = 3 principal components contributing the most to its total positional variance^62^. We interpret these components as the main linear motions explaining the collection’s conformational diversity. Namely, the *k*th principal component defines a set of 3D displacement vectors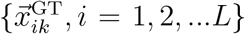 for the *L* protein residues’ C*α* atoms. We normalised these vectors to facilitate their comparison across different proteins, such that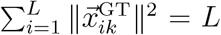. We further applied three filtering criteria with the aim of excluding collections with low diversity or highly non-linear complex deformations: (*i*) maximum Root Mean Squared Deviation (RMSD) between any two conformations of at least 2 °A, (*ii*) first principal component (main linear motion) contributing at least 80% of the total variance and (*iii*) involving at least 12 residues, *i*.*e*., *L× κ* ≥12, where *κ* is the collectivity of the principal component (see definition below). This operation resulted in 7 335 collections, randomly split between train (70%), validation (15%) and test (15%) sets.

In addition, we manually selected the conformational collections corresponding to the proteins from the iMod benchmark^68^ and put them in our test set. The iMod benchmark is derived from the molecular motions database MolMovDB^80^ and was previously used to assess coarse-grained elastic network model-based flexible fitting methods^81^. It comprises pairs of open-closed conformations for a couple of tens of proteins that represent a wide variety of motions, predominantly hinge motions but also shear and other complex motions. We could identify 21 conformational collections containing both open and closed conformations and complying with all our criteria, except for the percentage of variance explained by the first principal component (which can go down to 60% on this benchmark set).

DANCE makes use of a reference conformation to superimpose the C*α* atoms’ 3D coordinates and centre them prior to extracting motions with PCA. By default, the reference corresponds to the protein chain with the most representative amino acid sequence^62^. In order to augment the data, we defined up to 4 alternative reference conformations, in addition to the default one (option -n 5). At each iteration, DANCE chose the new reference conformation as the one displaying the highest RMSD from the previous one. This strategy maximises the impact of changing the reference and thus the diversity of the extracted motions.

#### Model Specifications

##### Input features

SeaMoon takes as input embeddings computed from pre-trained pLMs, namely Evolutionary Scale Models ESM2-T33-650M-UR50^23^ and ESM3-small (1.4B)^63^, as well as Protein sequence-structure T5^64^. ESM2-T33-650M-UR50 is a BERT^82^ style 650-million-parameter encoder-only transformer architecture trained on all clusters from Uniref50^83,84^, a version of UniProt^85^ clustered at 50% sequence similarity, augmented by sampling sequences from the Uniref90 clusters of the representative chains (excluding artificial sequences). ESM3-small (1.4B) is a transformer-based^86^ all-to-all generative architecture that both conditions on and generates a variety of different tracks representing protein sequence, secondary and tertiary structure, solvent accessibility and function. It was trained on over 2.5 billion natural proteins collected from sequence and structure databases, including UniRef, MGnify^87^, OAS^88^ and the PDB^49^, augmented with synthetic sequences generated by an inverse folding model^63^. Protein sequence-structure T5 is a bilingual pLM trained on a high-quality clustered version of the AlphaFold Protein Structure Database^89,90^ to translate 1D sequences of amino acids into 1D sequences of 3Di tokens representing 3D structural states^91^ and vice versa. The 3Di alphabet, introduced by the 3D-alignment method Foldseek^91^, describes tertiary contacts between protein residues and their nearest neighbours. This 1D discretised representation of 3D structures is sensitive to fold change but robust to conformational rearrangements. Protein sequence-structure T5 expands on ProtT5-XL-U50^22^, an encoder-decoder transformer architecture^92^ trained on reconstructing corrupted amino acids from the Big Fantastic Database^93^ and UniRef50. Throughout the text, we refer to these pLMs as ESM2, ESM3 and ProstT5, respectively. We used the pre-trained pLMs as is, without fine-tuning their weights, and we gave them only amino acid sequences as input.

##### Model’s architecture

SeaMoon’s architecture is a convolutional neural network^94^ taking as input a sequence embedding of dimensions *L × d*, with *L* the number of protein residues and *d* the representation dimension of the chosen pLM, namely 1 280 for ESM2, 1 536 for ESM3, and 1 024 for ProstT5, and outputting *K* predicted tensors of dimensions *L ×* 3. It comprises a linear layer followed by two hidden 1-dimensional convolutional layers with filter sizes of 15 and 31, respectively, and finally *K* parallel linear layers (**Table S1**). SeaMoon’s convolutional architecture allows handling sequences of any arbitrary length *L* and preserving this dimension throughout the network. All layers were linked through the LeakyReLu activation function^95^, as well as 80% dropout^96^. We experimented with other types of architectures, including those based on sequence transformers, and chose the one based on CNNs as it demonstrated the maximum accuracy at a reasonable number of trained parameters. Please see **Table S4** and **Fig. 9** for more details. We implemented the models in PyTorch^97^ v2.1.0 using Python 3.11.9.

By design, the SeaMoon model predicts the *K* motion tensors in a latent space that is invariant to the protein’s actual 3D orientation. To align these predictions with a given 3D conformation, additional information, such as the ground-truth motions, is required, as explained below.

##### Loss function

We aim to minimise the discrepancy between the predicted tensor *X* and the ground-truth tensor *X*^GT^, both of dimensions *L× K ×*3, expressed as a weighted aligned sum-of-squares error loss,

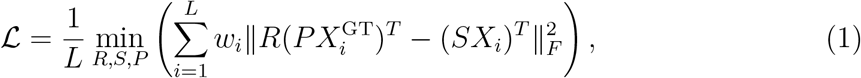

where *X*_*i*_ defines the set of *K* 3D displacements vectors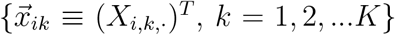predicted for the C*α* atom of residue *i*,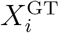 defines the corresponding ground-truth 3D displacement vector set‖·‖, _*F*_ designates the Frobenius norm, and *w*_*i*_ is a weight reflecting the confidence in the ground-truth data for residue *i*^62^. It is computed as the proportion of conformations in the experimental collection with resolved 3D coordinates for residue *i*.The matrices *R*, of dimension 3 *×*3, and *P*, of dimension *K ×K*, allow for rotating and permuting the ground-truth vectors to optimally align them with the predicted ones. We chose to apply the transformations to the ground-truth vectors for gradient stability. We allow for rotations *R* because SeaMoon relies solely on a protein sequence embedding as input. Its predictions are not anchored in a particular 3D structure and hence, they may be in any arbitrary orientation. We allow for permutation *P* to stimulate knowledge transfer across conformational collections. The rationale is that a motion may be shared between two collections without necessarily contributing to their positional variance to the same extent. Additionally, we allow for scaling predictions with the *K × K* diagonal matrix *S*, so that SeaMoon can focus on predicting only the relative motion amplitudes between the amino acid residues.

In practice, we first jointly determine the optimal permutation *P* and rotation *R* of the ground-truth 3D vectors. We test all possible permutations, and, for each, we determine the best rotation by solving the orthogonal Procrustes problem^98,99^. We shall note that the optimal solution may be a pseudo-rotation, *i*.*e*., det(*R*) = *−* 1, which corresponds to the combination of a rotation and an inversion. The loss can then be reformulated as,

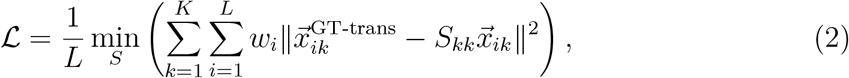

where 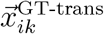 is the ground-truth 3D displacement vector for residue *i* matching the predicted 3D vector 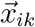 and aligned with it, and *S*_*kk*_ ∈ ℝ is the *k*th scaling coefficient, *i*.*e*. the *k*th non-null term of the diagonal scaling matrix *S*. The optimal value for *S*_*kk*_ is computed as,

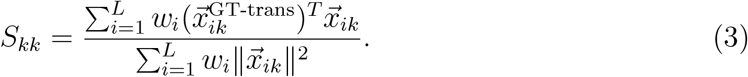

##### Training

We trained six models (**Table S2**) to predict *K* = 3 motions using the Adam optimizer^100^ with a learning rate of 1e-02. We used a batch size of 64 input sequences and employed padding to accommodate sequences of variable sizes in the same batch. We trained for 500 epochs and kept the best model according to the performance on the validation set.

##### Inference

We provide an unsupervised procedure to orient SeaMoon’s predicted motions with respect to a target 3D conformation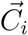during inference. This approach relies on the assumption that correct predictions comply with the same rotational constraints as ground-truth motions (see *Supplementary Methods*). Specifically, these constraints state that the cross products between the positional 3D vectors of the reference conformation *C*^0^ and the 3D displacement vectors defined by a ground-truth principal component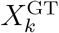result in a null vector,

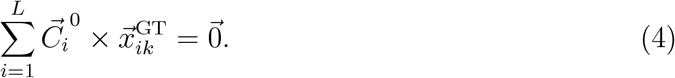

Assuming that the motion tensor *X*_*k*_ predicted by SeaMoon preserves this property, we determine the rotation *R* that minimises the following cross-product,

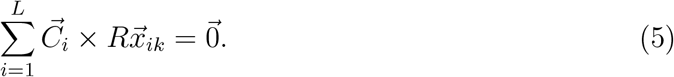

This problem has at most four solutions and we solve it exactly using the symbolic package *wolframclient* in Python. See *Supplementary Methods* for a detailed explanation. In practice, we observe that these four solutions reduce to two pairs of highly similar rotations.

##### Evaluation

We assessed SeaMoon predictions on each test protein from two different perspectives. In the first assessment, we considered all *K × K* pairs of predicted and ground-truth motions and estimated the discrepancy between the two motions within each pair after optimally rotating and scaling them. We focused on the best matching pair for computing success rates and illustrating the results. In the second assessment, we considered the predicted and ground-truth motion subspaces at once and estimated their permutation-, rotation- and scaling-invariant global similarity. In addition, we estimated discrepancies and similarities between individual predicted and ground-truth motions after globally matching and aligning the subspaces. We detail our evaluation metrics and procedures in the following.

##### Normalised sum-of-squares error

At inference time, we estimate the discrepancy between the *k*th predicted motion and the *l*th ground-truth principal component by computing their weighted sum-of-squares error under optimal rotation *R*^*opt*^ and scaling *s*^*opt*^,

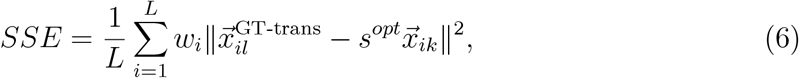

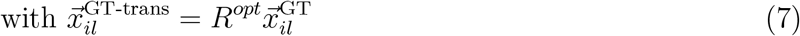

In the best-case scenario, the prediction is colinear to the transformed ground-truth,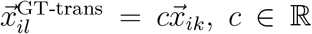, such that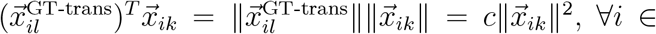 1, 2, …*L*. By virtue of 3, the scaling coefficient *s*^*opt*^ will be equal to *c*, and thus, the error will be null,

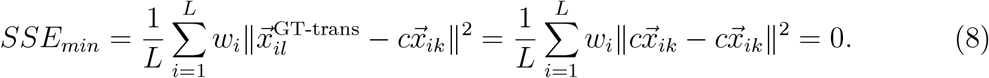

In the worst-case scenario, the prediction is orthogonal to the ground truth, such that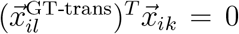, ∀*i* ∈ 1, 2, …*L*. The scaling coefficient will be null and, hence, this situation is equivalent to having a null prediction,

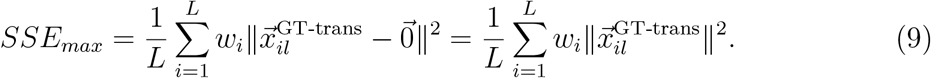

The value of the raw error depends on the uncertainty of the ground-truth data. If all conformations in the collection have resolved 3D coordinates for all protein residues, then *w*_*i*_ = 1, ∀*i* = 1, 2, …, *L* and the maximum error is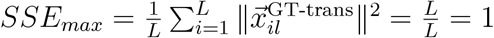As uncertainty in the ground-truth data increases, the associated errors will become smaller. To ensure a fair assessment of the predictions across proteins, we normalise the raw errors,

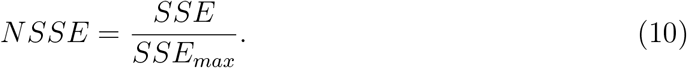

##### Subspace comparison

We estimated the similarity between the *K ×* 3 subspaces spanned by SeaMoon predictions and the ground-truth principal components as their Root Mean Square Inner Product (RMSIP)^73–75^. It is computed as an average of the normalised inner products of all the vectors in both subspaces,

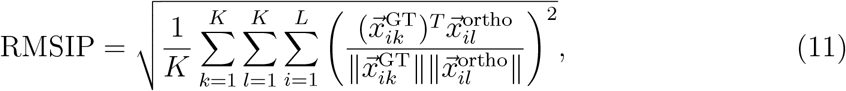

where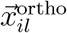is obtained by orthogonalising SeaMoon predictions using the Gram–Schmidt process. This operation ensures that the RMSIP ranges from zero for mutually orthogonalising subspaces to one for identical subspaces and avoids artificially inflating the RMSIP due to redundancy in the predicted motions. We should stress that in practice, this redundancy is limited and the motions predicted for a given protein never collapse (**Fig. S10**). A RMSIP score of 0.70 is considered an excellent correspondence while a score of 0.50 is considered fair^73^.

While the RMSIP is invariant to permutations and rotations, the individual inner products, reflecting similarities between pairs of motions, are not. For interpretability purposes, we maximised these pairwise similarities through the following procedure:

1. compute the NSSE for all pairs of predictions and ground-truth principal components, under optimal rotation and scaling, as in 7,
2. orthogonalise the predictions in the order of their losses, from the best-matching prediction to the worst-matching one,
3. determine the optimal global rotation of the ordered set of matching ground-truth components onto the ordered set of orthogonalised predictions,
4. compute all pairwise normalised inner products and the corresponding RMSIP, and all pairwise NSSE under optimal scaling.

### Comparison with the normal mode analysis

We compared SeaMoon performance with the physics-based unsupervised normal mode analysis (NMA) ^45^. The NMA takes as input a protein 3D structure and builds an elastic network model where the nodes represent the atoms and the edges represent springs linking atoms located close to each other in 3D space. The normal modes are obtained by diagonalizing the mass-weighted Hessian matrix of the potential energy of this network. We used the highly efficient NOLB method^47^ to extract the first *K* = 3 normal modes from the test protein 3D conformations. We retained only the C*α* atoms, as for the principal component analysis, and defined the edges in the elastic network using a distance cutoff of 10°A. We enhanced the elastic network dynamical potential by excluding edges corresponding to small contact areas between protein segments. We detected them as disconnected patches in the contact map using HOPMA^48^. Contrary to SeaMoon predictions, the orientation of the NMA predictions is not arbitrary and thus, we do not need to align the ground-truth components onto them.

### Protein fold-centered analysis

We defined protein folds as the topologies from the CATH classification^77^. We estimated the NSSE of each fold as the average NSSE computed over all test proteins containing the corresponding topology. At the protein level, we consider the NSSE of the best matching pair of predicted and ground-truth motions.

### Protein properties

#### Sequence and structure similarity

We estimated sequence similarity between train and test proteins using MMseqs2^101^ with default settings. We used TM-align (version 20220412) to perform all-to-all pairwise structural alignments between train and test protein conformations and compute TM-scores^66^. TM-score measures the topological similarity of protein structures. It ranges between 0 and 1, and a score higher than 0.5 assumes roughly the same fold. Furthermore, to account for protein flexibility in our estimation of structural similarity, we used the flexible structural alignment functionality from Kpax (version 5.1.1)^67,102^. Kpax implements a tiled dynamic programming algorithm that optimally partitions the input structures into rigid segments and aligns them. This strategy makes the TM-score estimation robust to changes in the relative positions and orientations of the identified segments, and thus to conformational changes. For each test protein, we computed the flexible Kpax TM-score with its best training hit identified by TM-align.

#### Motion contribution and collectivity

We estimate the contribution of the *L×*3 of the ground-truth principal component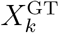to the total positional variance as its normalised eigenvalue, 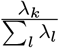. We estimate the collectivity^103,104^ motion tensor *X*_*k*_ as,

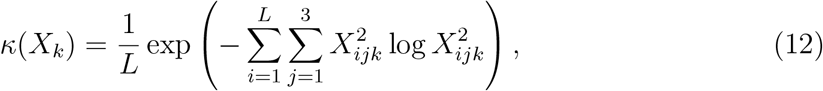

with *L* the number of residues. If *κ*(**v**) = 1, then the corresponding motion is maximally collective and has all the atomic displacements identical. In case of an extremely localised motion, where only one single atom is affected, the collectivity is minimal and equals to 1*/L*.

#### Visualisation

Three-dimensional protein structures were visualised and rendered using PyMOL^105^. All plots and statistical visualisations were generated using the ggplot2 R package version 4.2^106^. Motion directions were represented as 3D arrows overlaid on the structures using a custom Python script developed in-house that utilises PyMOL’s CGO module and NumPy^107^ for vector calculations.

### Quantification and statistical analysis

We assessed the statistical significance of performance differences between ESM3 and ProstT5 with a paired Wilcoxon signed-rank test^108,109^ performed using the *wilcox*.*test* function implemented in the R Stats Package version 4.2.2^110^. We assessed the statistical significance of the enrichments and depletions in motion types reported in Figure 3A with hypergeometric tests using the *phyper* function from the same package. Furthermore, we quantified the significance and statistical reliability of SeaMoon performance by comparing it with a random baseline and by performing bootstrap resampling, as described below.

#### Estimation of sum-of-squares errors for random vectors

To compare SeaMoon results with a random baseline, we selected 14 ground-truth principal components from the test set. We focused on proteins with maximum confidence, *i*.*e*., for which *w*_*i*_ = 1, ∀*i* = 1, 2, …, *L*. We started with a set of 10 components chosen randomly. We then added the most localised component (collectivity *κ* = 0.06), the most collective one (*κ* = 0.85), a component from the smallest protein (33 residues), and a component from the longest one (662 residues). We generated 1000 random predictions for each ground truth component and computed their sum-of-squares errors under optimal rotation and scaling.

#### Success rate estimation

We estimated the success rate of a method as the percentage of test proteins for which it approximated at least one ground-truth motion with normalised sum-of-squares error NSSE*<*NSSE_*cut*_, where NSSE_*cut*_ is set to 0.6 by default. To assess the statistical reliability of our success rate estimates, we used bootstrap resampling. Specifically, we generated 1 000 bootstrap random samples of *N* ^2*/*3^ test proteins, where *N* =1 121 is the cardinality of the full test set, allowing for replacement. For each method and each NSSE cutoff considered, we computed the average success rate, as well as the interquartile range and 95% central range over the 1 000 bootstrap samples.

### Key resources table

| REAGENT or RESOURCE | SOURCE | IDENTIFIER | ADDITIONAL INFORMATION |
| --- | --- | --- | --- |
| **Deposited data** |  |  |  |
| Protein motion dataset | This paper | 10.5281/zenodo.13833309 | https://zenodo.org/records/13833309 |
| SeaMoon trained models | This paper | 10.5281/zenodo.15616637 | https://github.com/PhyloSofS-Team/seamoon |
| **Software and algorithms** |  |  |  |
| Python | Python Software Foundation | RRID:SCR_008394 | Version 3.8+ |
| PyTorch | 97 | RRID:SCR_018536 | Version 2.1.0 |
| PyMOL | 105 | RRID:SCR_000305 | Version 2.5 |
| R | R Core Team | RRID:SCR_001905 | Version 4.2.2 |
| ggplot2 | 106 | RRID:SCR_014601 | Version 4.2 |
| TM-align | 66 | N/A | Version 20220412 |
| Kpax | 67,102 | N/A | Version 5.1.1 |
| ESM-2 | 23 | N/A | https://huggingface.co/facebook/esm2_t33_650M_UR50D |
| ESM-3 | 63 | N/A | https://huggingface.co/EvolutionaryScale/esm3-sm-open-v1 |
| ProstT5 | 64 | N/A | https://huggingface.co/Rostlab/ProstT5 |
| DANCE | 62 | 10.5281/zenodo.15617758 | https://github.com/PhyloSofS-Team/DANCE |
| HOPMA | 48 | 10.5281/zenodo.15616796 | https://github.com/elolaine/HOPMA |
| NOLB | 47 | N/A | https://team.inria.fr/nano-d/software/nolb-normal-modes/ |
| SeaMoon code | This paper | 10.5281/zenodo.15616637 | https://github.com/PhyloSofS-Team/seamoon |
| **Other** |  |  |  |
| Protein Data Bank | wwPDB | RRID:SCR_012820 | https://www.rcsb.org/ |
| PDB-redo | 79 | N/A | https://pdb-redo.eu |

## Notes

### Competing Interest Statement

The authors have declared no competing interest.

### Summary of Updates

- results: additional calculations to strengthen the statistical significance of the results, clarification of the choice of thresholds - introduction and discussion: add sentences about the biological relevance of the work. - some figures were moved from supp to main.

