## supplemental material for "SeaMoon: from protein language models to continuous structural heterogeneity"

### Supplemental methods

We compute the ground-truth motions with DANCE [?] by applying Principal Component Analysis on the 3D coordinates of a collection of protein conformations superimposed onto one another. The superimposition puts the protein conformations' centers of mass to zero and then aims at determining the optimal least-squares rotation matrix minimizing the Root Mean Square Deviation (RMSD) between any conformation and a reference conformation. Let us first derive some preliminary properties of this optimal rigid superposition.

#### Implications of optimal superimposition

Let us consider two sets of 3-vectors $\mathcal{A} = \{\vec{a}_i, i = 1, ...., m\}$ and $\mathcal{B} = \{\vec{b}_i, i = 1, ...., m\}$, such that a rigid transformation defined by the translation vector $\vec{T}$ and the rotation matrix $R$ minimizes their squared geometric mismatch:

$$\sum_i (\vec{a}_i - R\vec{b}_i - \vec{T})^2 \to \min . \tag{1}$$

By taking the derivative of the above equation with respect to the rigid translation $\vec{T}$, we obtain

$$\sum_i (\vec{a}_i - R\vec{b}_i - \vec{T}) = \vec{0}. \tag{2}$$

Similarly, by taking the derivative of the above equation with respect to the rotation matrix $R$ about $x$, $y$, and $z$ axes, we obtain

$$\sum_i \vec{b}_i \times (\vec{a}_i - R\vec{b}_i - \vec{T}) = \vec{0}, \tag{3}$$

where $\vec{b}_i \times (\vec{a}_i - R\vec{b}_i - \vec{T})$ is the angular velocity of the particle $i$ from $\mathcal{B}$. Therefore, the total translational force and the total rotational force or torque applied to a set of points in a rigid body must be zero at the optimal superposition conformation.

#### Ground-truth motions' translational and rotational constraints

The Cartesian coordinates of the 3D protein conformations in a collection are stored in a matrix $X$ of dimension $3m \times n$. Their positional covariance matrix $C$ of dimensions $3m \times 3m$ is expressed as,

$$C = \frac{1}{n-1} X X^T. \tag{4}$$

The eigen decomposition of the covariance matrix $C = VDV^T$ leads to a set of eigenvectors $\{V_{\cdot j},\ j = 1, ..., 3m\}$ which are the columns of the matrix $V$, associated with the set of eigenvalues $\{\lambda_j,\ j = 1, ..., 3m\}$ stored in the diagonal matrix $D$. We show below that these eigenvectors comply with specific translational and rotational constraints.

**Translational constraint.** Since the conformations are centred at the origin, we have $\sum_{l=1}^{m} \vec{x}_{lj} = \vec{0}$ for every conformation $j = 1, ..., n$, with $\vec{x}_{lj}$ the 3-component vector from $X$ corresponding to atom $l$. It follows that the elements of each $x$,$y$,$z$ component in each ow of the covariance matrix $C$ sum to zero,

$$\sum_{x_j,y_j,z_j=1}^{m} C_{ij} = \frac{1}{n-1} \sum_{x_j,y_j,z_j=1}^{m} \sum_{k=1}^{n} X_{ik} X_{jk} = \frac{1}{n-1} \sum_{k=1}^{n} X_{ik} \sum_{x_j,y_j,z_j=1}^{m} X_{jk} = 0. \tag{5}$$

Without loss of generality, we can define coordinate indexes as $j_x = 3j$, $j_y = 3j + 1$, and $j_z = 3j + 2$. According to the eigen decomposition, we can then write, $\forall i$, $i = 1, ..., 3m$,

$$\sum_{x_j,y_j,z_j=1}^{m} C_{ij} = \sum_{x_j,y_j,z_j=1}^{m} \sum_{k=1}^{3m} \lambda_k V_{ik} V_{jk} = 0. \tag{6}$$

Summing up over the $x$,$y$, and $z$ components of the rows of $C$, we obtain,

$$\sum_{x_i,y_i,z_i=1}^{m} \sum_{x_j,y_j,z_j=1}^{m} C_{ij} = \sum_{x_i,y_i,z_i=1}^{m} \sum_{x_j,y_j,z_j=1}^{m} \sum_{k=1}^{3m} \lambda_k V_{ik} V_{jk} = 0, \tag{7}$$

which can be rewritten as,

$$\lambda_1 \left(\sum_i V_{i1}^x\right)^2 + \lambda_2 \left(\sum_i V_{i1}^y\right)^2 + \lambda_3 \left(\sum_i V_{i1}^z\right)^2 + \lambda_4 \left(\sum_i V_{i2}^x\right)^2 \lambda_5 \left(\sum_i V_{i2}^y\right)^2 + \lambda_6 \left(\sum_i V_{i2}^z\right)^2 \tag{8}$$

$$+ ... + \lambda_{3m-2} \left(\sum_i V_{i(m)}^x\right)^2 + \lambda_{3m-1} \left(\sum_i V_{i(m)}^y\right)^2 + \lambda_{3m} \left(\sum_i V_{i(m)}^z\right)^2 = 0. \tag{9}$$

Since the covariance matrix $C$ is symmetric and positive semi-definite, all the eigenvalues $\lambda_k$ are real and non-negative and it follows that the sum of components of each eigenvector $V_{\cdot k}$ corresponding to a positive eigenvalue $\lambda_k$ must be zero. And the number of such eigenvectors equals the rank of the covariance matrix.

**Rotational constraint.** We can express the angular velocity of an atom $l$ from conformation $j$ from our set as $\vec{r}_{lj} \times \vec{x}_{lj}$, where $\vec{r}_{lj}$ is its 3D position relative to the conformation's center of mass, *i.e.*, the rotation center, and $\vec{x}_{lj}$ is its 3D displacement vector from the reference conformation. This angular velocity can be re-written as $(\vec{r}_l + \vec{x}_{lj}) \times \vec{x}_{lj} = \vec{r}_l \times \vec{x}_{lj}$, where $\vec{r}_l$ is the 3D position of the atom $l$ in the *reference* conformation relative its center of mass. We can further rewrite this vector product in a matrix form as $[r_l]_\times \vec{x}_{lj}$, where $[r_l]_\times$ is a $3 \times 3$ skew-symmetric matrix that corresponds to the cross product operation. Let us also define a $3m \times 3m$ block-diagonal matrix $R$ formed of $m$ matrices $[r_i]_\times$, $R \equiv \text{diag}([r_1]_\times, [r_2]_\times, ..., [r_m]_\times)$.

Since the global rotations between protein conformations, with respect to the reference, have been removed during superimposition, the angular velocities for any conformation $j$ result in a null vector, $\sum_{l=1}^{m} [r_l]_\times \vec{x}_{lj} = \vec{0}$. Or, in the matrix form, for each column (conformation) $j$, $\sum_{i=1}^{m} (RX)_{i_x j} = 0$, $\sum_{i=1}^{m} (RX)_{i_y j} = 0$, $\sum_{i=1}^{m} (RX)_{i_z j} = 0$, where, without loss of generality, we can define coordinate indexes as $i_x = 3i$, $i_y = 3i+1$, and $i_z = 3i + 2$. It follows that,

$$\forall a \in \{x, y, z\}: \quad \sum_{j=1}^{m} (RX(RX)^T)_{ij_a} = \sum_{j=1}^{m} \sum_{k=1}^{n} (RX)_{ik} (RX)_{j_a k} = \sum_{k=1}^{n} (RX)_{ik} \sum_{j=1}^{m} (RX)_{j_a k} = 0. \tag{10}$$

The decomposition of $RX(RX)^T$ leads to, $\forall i$, $i = 1, ..., 3m$,

$$\forall a \in \{x, y, z\}: \quad \sum_{j=1}^{m} (RX(RX)^T)_{ij_a} = \sum_{j=1}^{m} (R[\sum_k \lambda_k V_k V_k^T] R^T)_{ij_a} = \sum_k \lambda_k \sum_{j=1}^{m} (RV_k [RV_k]^T)_{ij_a} = 0 \tag{11}$$

Summing up over the $x$,$y$, and $z$ components of the rows of the previous system, we obtain,

$$\lambda_1 \left(\sum_i (RV_1)_{i_x}\right)^2 + \lambda_1 \left(\sum_i (RV_1)_{i_y}\right)^2 + \lambda_1 \left(\sum_i (RV_1)_{i_z}\right)^2 + \tag{12}$$

$$\lambda_2 \left(\sum_i (RV_2)_{i_x}\right)^2 + \lambda_2 \left(\sum_i (RV_2)_{i_y}\right)^2 + \lambda_2 \left(\sum_i (RV_2)_{i_z}\right)^2 + \tag{13}$$

$$... + \lambda_{3m} \left(\sum_i (RV_{3m})_{i_x}\right)^2 + \lambda_{3m} \left(\sum_i (RV_{3m})_{i_y}\right)^2 + \lambda_{3m} \left(\sum_i (RV_{3m})_{i_z}\right)^2 = 0. \tag{14}$$

Thus, the positional eigenvectors must have additional rotational constraints, $\forall k$, $k = 1, ..., 3m$, $\sum_i (RV_k)_{i_x} = 0$ ,$\sum_i (RV_k)_{i_y} = 0$ , and $\sum_i (RV_k)_{i_z} = 0$ for all eigenvectors with the corresponding non-zero eigenvalues.

### Orienting a predicted motion with respect to a 3D conformation

We exploit the rotational constraints of the ground-truth motions, *i.e.*, the eigenvectors of the positional covariance matrix, to align the motion vectors predicted by SeaMoon on a given protein 3D conformation. More specifically, we aim at determining the rotation $R \in \mathrm{SO}(3)$ that minimizes the overall angular velocity of the conformation subjected to the predicted motion. If we denote $(v^i)_{i \leq m}$ the 3D displacement vectors predicted by SeaMoon for $m$ protein atoms, the problem we solve is $\sum_i (Rv^i) \times r^i = 0$ for the rotation $R$, where $r^i$ is the 3D positional vector of atom $i$.

For coordinate $d \in \{1, 2, 3\}$, we can then write, using Einstein's notation:

$$
\begin{aligned}
0 &= e_d.(Rv^i) \times r^i \\
&= \varepsilon_{pqd}(Rv^i)_p r^i_q \\
&= \varepsilon_{pqd} R_{pk} v^i_k r^i_q \\
&= R_{pk} \varepsilon_{pqd} v^i_k r^i_q \\
&= \mathrm{Tr}(R A_d^T)
\end{aligned}
$$

where $(A_d)_{pk} = \varepsilon_{pqd} v^i_k r^i_q$ and where $\varepsilon_{i,j,k}$ is 0 if $i, j, k$ are not different and elsewise is equal to the signature of the permutation

$$
\begin{pmatrix} 1 & 2 & 3 \\ i & j & k \end{pmatrix}
$$

To solve these equations, we use 4-dimensional quaternions which have fewer dimensions than $3 \times 3$ matrices, associating to a unitary quaternion the rotation matrix [**?**]:

$$
R_q = \begin{pmatrix} q_1^2 + q_2^2 - q_3^2 + q_4^2 & 2(q_2q_3 + q_1q_4) & 2(q_2q_4 - q_1q_3) \\ 2(q_2q_3 - q_1q_4) & q_1^2 + q_3^2 - q_2^2 + q_4^2 & 2(q_3q_4 + q_1q_2) \\ 2(q_2q_4 + q_1q_3) & 2(q_3q_4 - q_1q_2) & q_1^2 + q_4^2 - q_2^2 + q_3^2 \end{pmatrix} \tag{15}
$$

We can develop the expression $\mathrm{Tr}(RA^T)$ to give:

$$
\begin{aligned}
\mathrm{Tr}(RA^T) = &(-2q_1q_2 + 2q_3q_4)A_{3,2} + (2q_1q_2 + 2q_3q_4)A_{2,3} \\
&+ (-2q_1q_3 + 2q_2q_4)A_{1,3} + (2q_1q_3 + 2q_2q_4)A_{3,1} \\
&+ (-2q_1q_4 + 2q_2q_3)A_{2,1} + (2q_1q_4 + 2q_2q_3)A_{1,2} \\
&+ (q_1^2 - q_2^2 - q_3^2 + q_4^2)A_{3,3} + (q_1^2 - q_2^2 + q_3^2 - q_4^2)A_{2,2} + (q_1^2 + q_2^2 - q_3^2 - q_4^2)A_{1,1}
\end{aligned}
$$

Writing the same equations changing $A$ for $A_m$ for $m = 1, 2, 3$, we get a system of 3 quadratic equations for 4 unknowns by setting the previous terms to 0, to which we add the unitary constraint $q_0^2 + q_1^2 + q_3^2 + q_4^2 = 1$. We solve this system using the symbolic package *wolframclient* in Python, yielding at most 4 quaternions as solutions which we then transform in rotation matrices.

### Supplemental tables and figures

| Layer type | Output shape | Filter size | #(Parameters) |
|---|---|---|---|
| Input | ESM2:($L$, 1 280)<br>ESM3:($L$, 1 536)<br>ProstT5:($L$, 1 024) | - | 0 |
| Conv1D | (256, $L$) | 1 | ESM2: 327 936<br>ESM3: 393 216<br>ProstT5: 262 400 |
| Conv1D | (128, $L$) | 15 | 491 648 |
| Conv1D | (64, $L$) | 31 | 254 016 |
| $K$ Conv1D | ($K$, 3, $L$) | 1 | $K \times 195$ |

Supplemental Table S1: **Description of SeaMoon neural network architecture.** We report the layer types, output shapes, kernel sizes, and parameters for ESM2-, ESM3-, and ProstT5-based SeaMoon models. The length (in amino acid) of the input sequence is denoted as $L$ and the number of predicted vectors as $K$. The 1D convolutional layers with filter size of 1 are equivalent to linear layers.

| Method | Input | pLM | Supervised | #(Train samples) |
|---|---|---|---|---|
| SeaMoon-ESM2 | sequence | ESM2 | ✓ | 5 119 |
| SeaMoon-ESM2(x5) | sequence | ESM2 | ✓ | 14 921 |
| SeaMoon-ESM3 | sequence | ESM3 | ✓ | 5 119 |
| SeaMoon-ESM3(x5) | sequence | ESM3 | ✓ | 14 921 |
| SeaMoon-ProstT5 | sequence | ProstT5 | ✓ | 5 119 |
| SeaMoon-ProstT5(x5) | sequence | ProstT5 | ✓ | 14 921 |
| NMA | 3D structure | × | × | 0 |

Supplemental Table S2: **Description of the tested models and methods.** For SeaMoon-ESM2(x5), SeaMoon-ESM3(x5) and SeaMoon-T5(x5), we increased the number of training samples by defining up to 5 reference conformations per experimental collection.

Supplemental Table S3: **Performance stratified by different motion sizes**

| **Method** | **Protocol** | **Number of motions** | **Proportion of motions (p-value)** | | |
|---|---|---|---|---|---|
| | | | local $\kappa \leq 0.30$ | regional $0.30 < \kappa \leq 0.60$ | global $\kappa > 0.60$ |
| Test set | | 3 363 | 0.37 | 0.42 | 0.21 |
| SeaMoon | ESM2 | 350 | 0.35 (-1.8e-01) | 0.32 (-4.9e-05) | 0.33 (1.7e-08) |
| | ESM2(x5) | 384 | 0.30 (-9.5e-04) | 0.36 (-7.9e-03) | 0.34 (1.7e-10) |
| | ESM3 | 449 | 0.36 (-2.5e-01) | 0.33 (-1.7e-05) | 0.32 (1.0e-08) |
| | ESM3(x5) | 476 | 0.32 (-4.2e-03) | 0.34 (-1.8e-04) | 0.34 (6.6e-13) |
| | ProstT5 | 486 | 0.30 (-2.2e-04) | 0.34 (-1.3e-04) | 0.36 (2.2e-16) |
| | ProstT5(x5) | 501 | 0.28 (-5.3e-07) | 0.36 (-1.7e-03) | 0.37 (8.4e-19) |
| NMA | | 381 | 0.08 (-1.0e-44) | 0.40 ( -2.7e-01) | 0.52 (2.8e-47) |

We report the proportions of local, regional and global motions in the whole test set (3 ground-truth motions for each of the 1 121 test proteins) and in the sets of acceptable predictions generated by each method. We computed the signed p-values indicated in parenthesis with a hypergeometric test – positive for enrichment, negative for depletion.

| **Ablation applied** | **# of proteins w. acceptable prediction** |
|---|---|
| Base model, SeaMoon-T5 | 439 (39.2%) |
| **Network architecture**: | |
| Size 1 kernel | 271 (24.2%) |
| 7-layer Transformer architecture | 411 (36.7%) |
| **Training loss**: | |
| Without sign flip | 413 (36.8%) |
| Without permutation | 375 (33.4%) |
| Without reflection | 402 (35.9%) |
| **Input data**: | |
| Random embeddings | 119 (10.6%) |
| Positional encoding only | 177 (15.8%) |
| **Random baseline**: | |
| Random neural network weights | 0 (0.0%) |

Supplemental Table S4: **Success rate in ablation study.** The transformer architecture comprises one linear layer from dimension 1024 to 128, seven Transformer layers of size 128, with 4 heads, and three 1D CNNs with kernel size 1 (same as the base model) going from dimension 128 to 3 for each predicted mode. It has a similar number of free parameters compared to the base model.

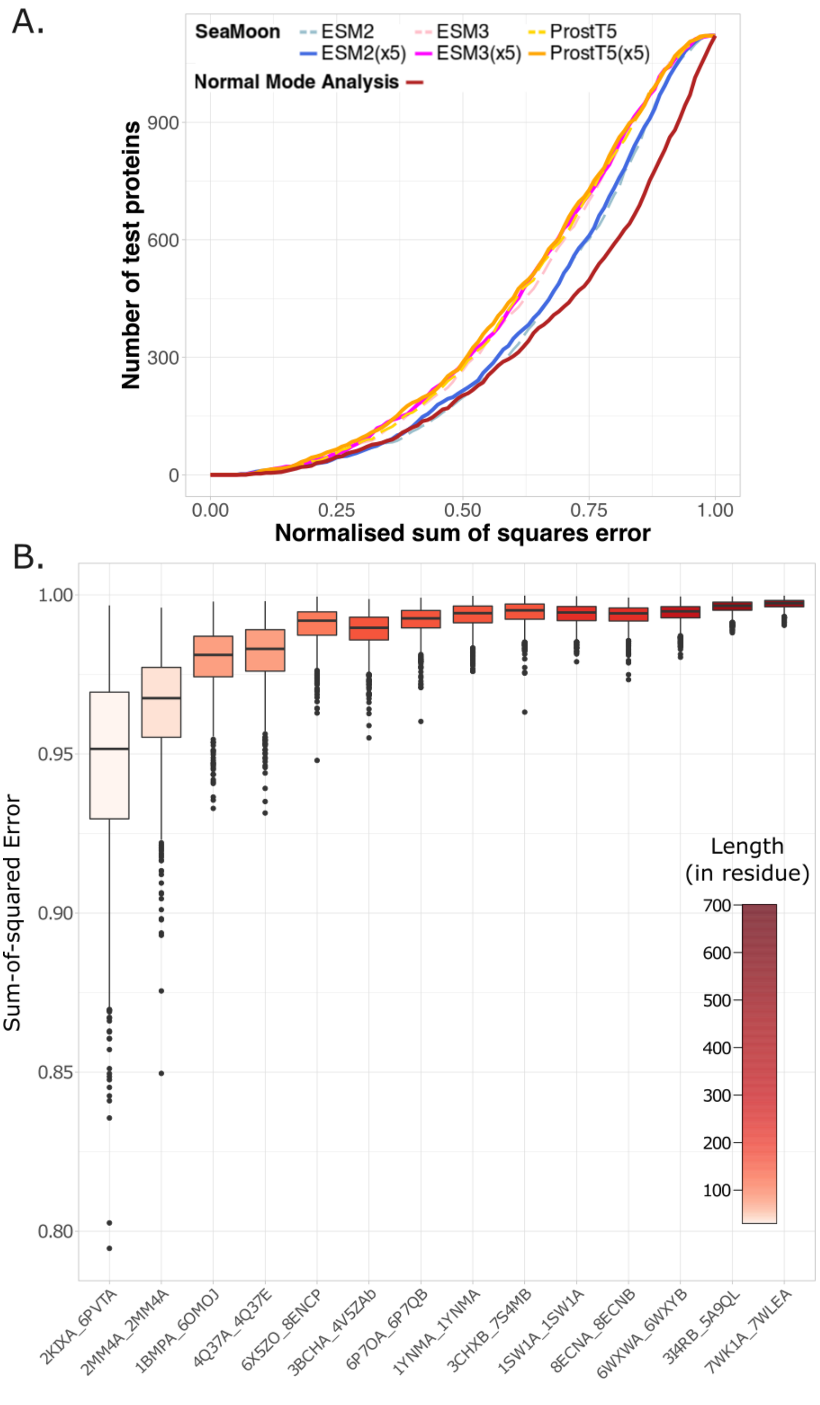


Supplemental Figure S1: **Performance on a test set of 1 121 proteins, related to Figure 2. A.** Comparison of the cumulative normalised sum-of-squares error (SSE) curves computed for different versions of SeaMoon and for Normal Mode Analysis (NMA, performed with NOLB). **B.** Distributions of normalised sum-of-squares errors computed after optimal rotation and scaling of 1000 random vectors against 14 ground-truth motions from the test set. The PDB chain identifiers of the corresponding proteins are given in x-axis. The boxes are colored according to the protein length.

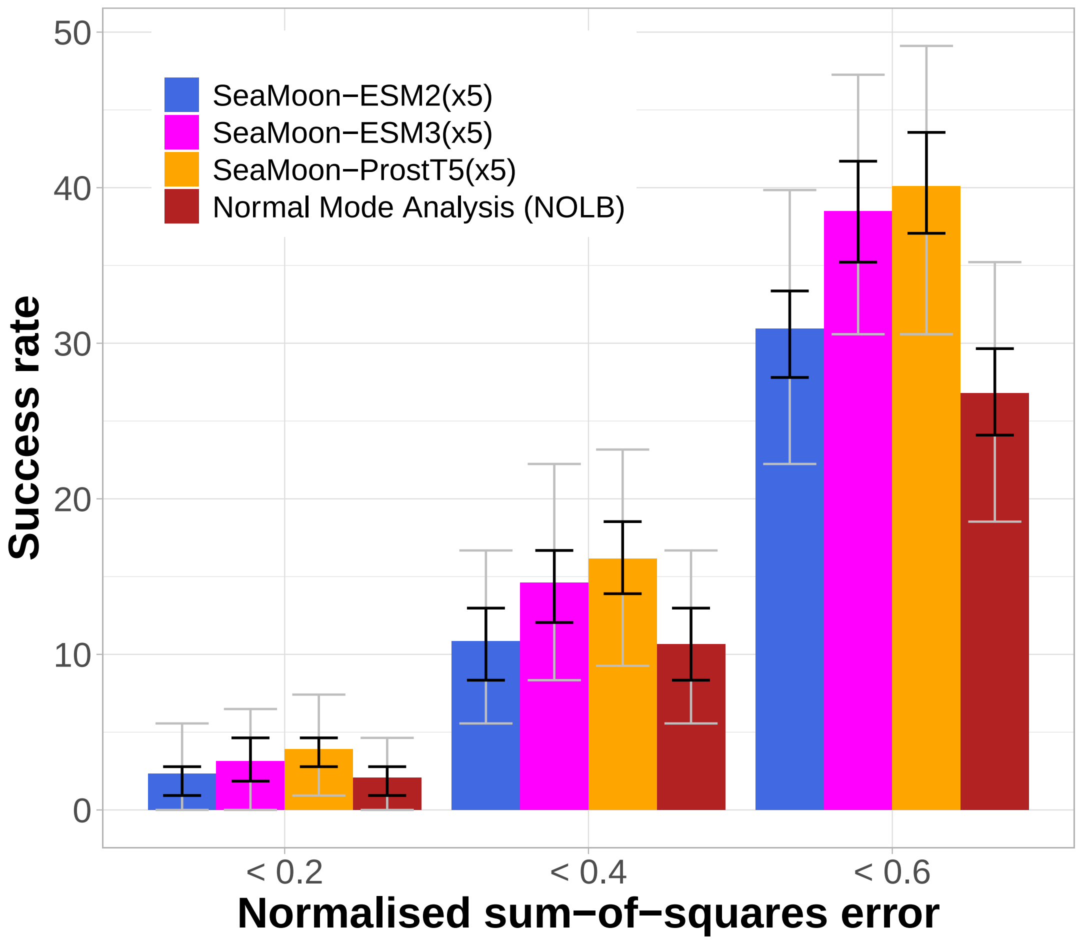


Supplemental Figure S2: **Success rate comparison and robustness, related to Table 1.** Average success rates computed over 1 000 bootstrap samples randomly drawn from the test set, with replacement. Each sample contains $N^{2/3} = 108$ proteins, where $N$=1 121 is the total number of test proteins. We estimate the success rate as the percentage of proteins for which at least one predicted motion approximates a ground-truth motion with NSSE below 0.2, 0.4, or 0.6, respectively. The black error bars indicate the interquartile ranges and the grey ones the 95% central ranges of the bootstrap estimates.

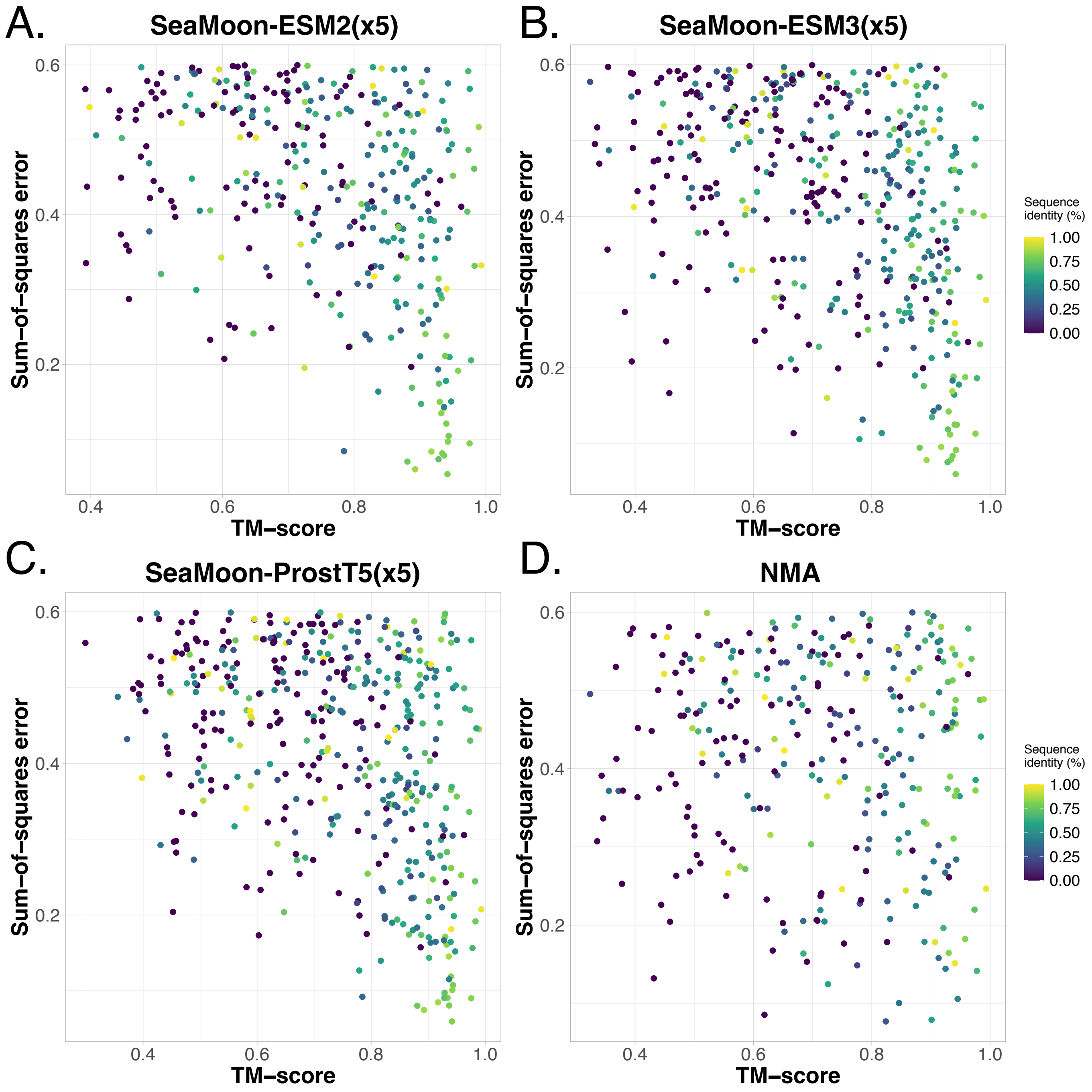


Supplemental Figure S3: **Influence of sequence and structure similarity, related to Figure 2.** Normalised sum-of-squares errors (y-axis) in function of the maximum TM-score (x-axis) and maximum sequence identity (color) of the test conformations computed over the whole training set. **A.** SeaMoon-ESM2(x5). **B.** SeaMoon-ESM3(x5). **C.** SeaMoon-ProstT5(x5). **D.** NMA.

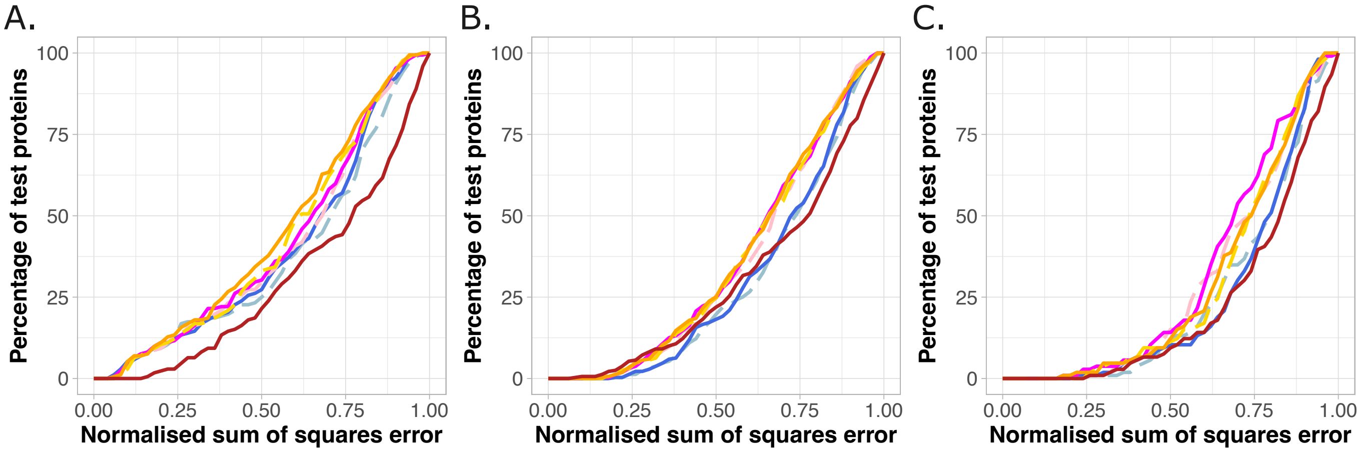


Supplemental Figure S4: **Comparison of performance depending on structural similarity, related to Figure 2.** Cumulative NSSE computed on three subsets of test proteins with increasing difficulty. **A. Easy:** at least 70% sequence identity with at least one train set example. **B. Intermediate:** at most 30% sequence identity and a TM-score of at least 0.7 with any train set example. **C. Difficult:** at most 30% sequence identity and 0.5 as TM-score with any train set example.The TM-score estimation was performed using the flexible structural alignment functionality of Kpax. It accounts for protein backbone flexibility. The three subsets contain 172, 319, and 106 proteins, respectively.

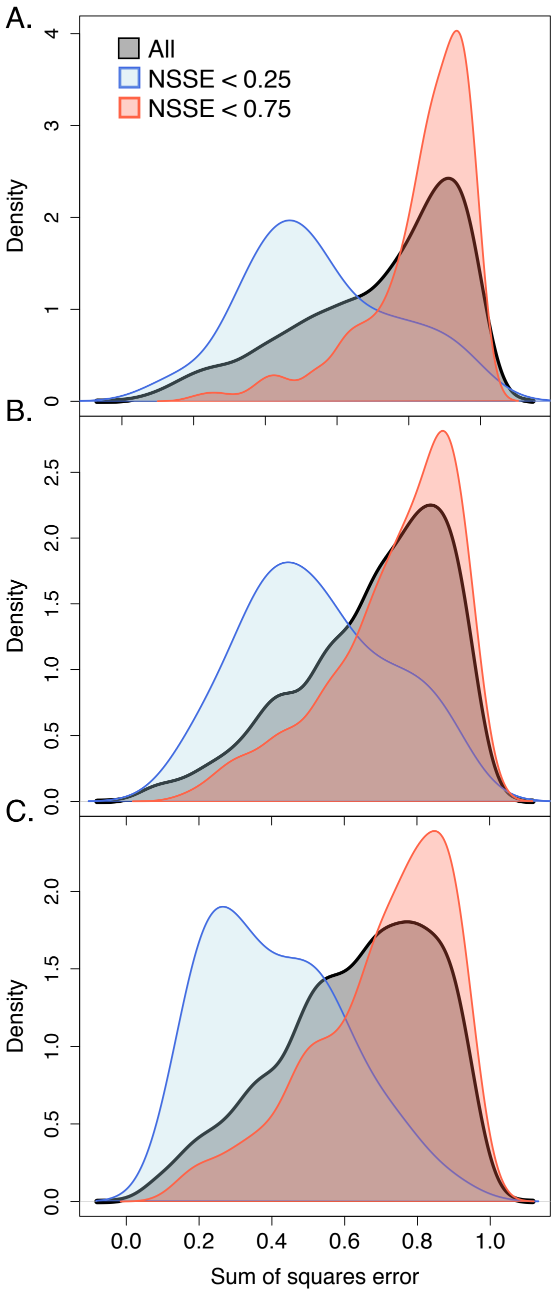


Supplemental Figure S5: **Agreement between a selection of methods, related to Figure 2. A.** Distribution densities of the NSSE computed for the NMA over the full test set (in black, 1 121 proteins) and over the subsets best-predicted ($NSSE < 0.25$, in blue, 36 proteins) and worst-predicted ($NSSE > 0.75$, in red, 326 proteins) by SeaMoon-ESM2(x5) and SeaMoon-ProstT5(x5). **B-C.** Distribution densities of the NSSE computed for SeaMoon-ESM2(x5) (B) and SeaMoon-ProstT5(x5) (C) over the full test set (in black) and over the subsets best-predicted ($NSSE < 0.25$, in blue, 46 proteins) and worst-predicted ($NSSE > 0.75$, in red, 624 proteins) by the NMA.

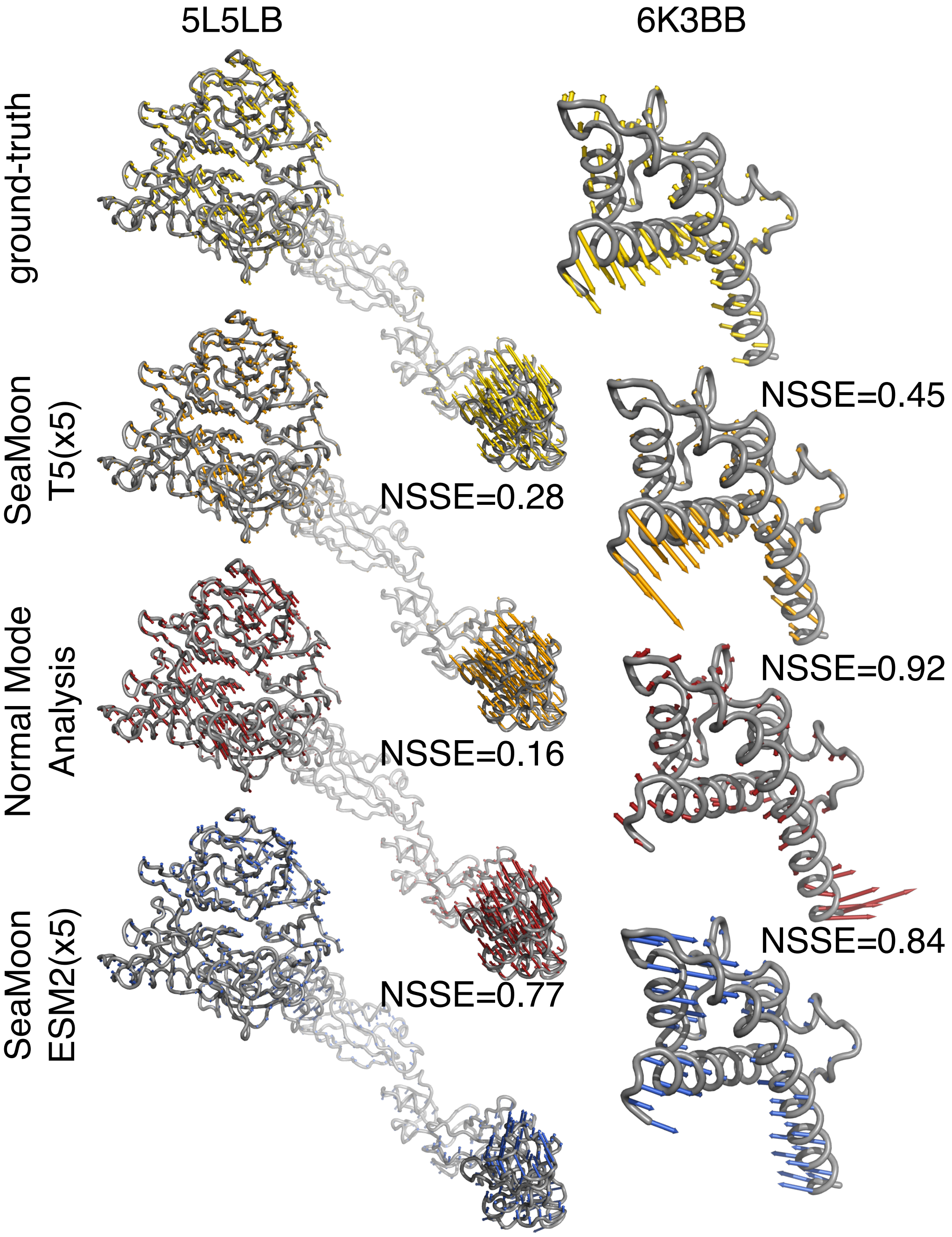


Supplemental Figure S6: **Examples of motions better captured by SeaMoon-ProstT5(x5) than SeaMoon-ESM2(x5), related to Figure 2.** The arrows depicted in yellow, orange, red, and blue onto the 3D structures represent the ground-truth motions and the best-matching predictions from SeaMoon-ProstT5(x5), the NMA, and SeaMoon-ESM2(x5), respectively. Left: Mammalian plexin A4 ectodomain (PDB code: 5L5L, chain B). It shares 64% sequence similarity with a plexin A2 ectodomain from the train set (TM-score = 0.68). Right: Legionella effector MavC (PDB code: 6K3B, chain B). It does not have any detectable sequence similarity with the training set and shares only a weak structural similarity (TM-score = 0.59) with the mammalian cytochrome P450 2B4.

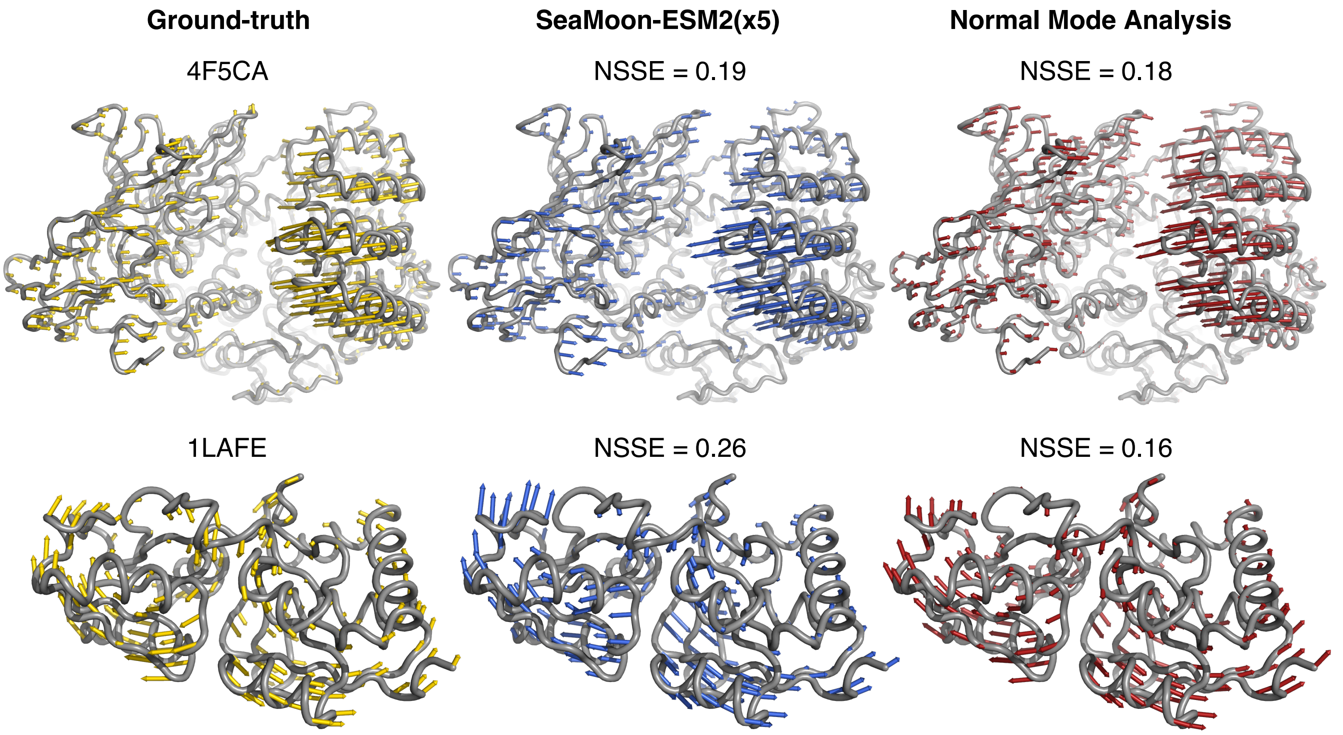


Supplemental Figure S7: **Examples of motions well predicted by SeaMoon-ESM2(x5) and the NMA, related to Figure 5.** The arrows depicted in yellow (left), blue (middle) and red (right) onto the 3D structure represent the ground-truth motion, the best-matching prediction from SeaMoon-ESM2(x5), and the best-matching prediction from the NMA. Top: Mammalian aminopeptidase N (PDB code: 4F5C, chain A). It shares 81% sequence similarity with a human aminopeptidase from the train set (TM-score = 0.96). Bottom: Bacterial periplasmic lysine-, arginine-, ornithine-binding protein (PDB code: 1LAF, chain E). It shares only 35% sequence similarity with its closest homolog from the train set, a nopaline-binding periplasmic protein from bacteria. Their structures are highly similar, with a TM-score of 0.91.

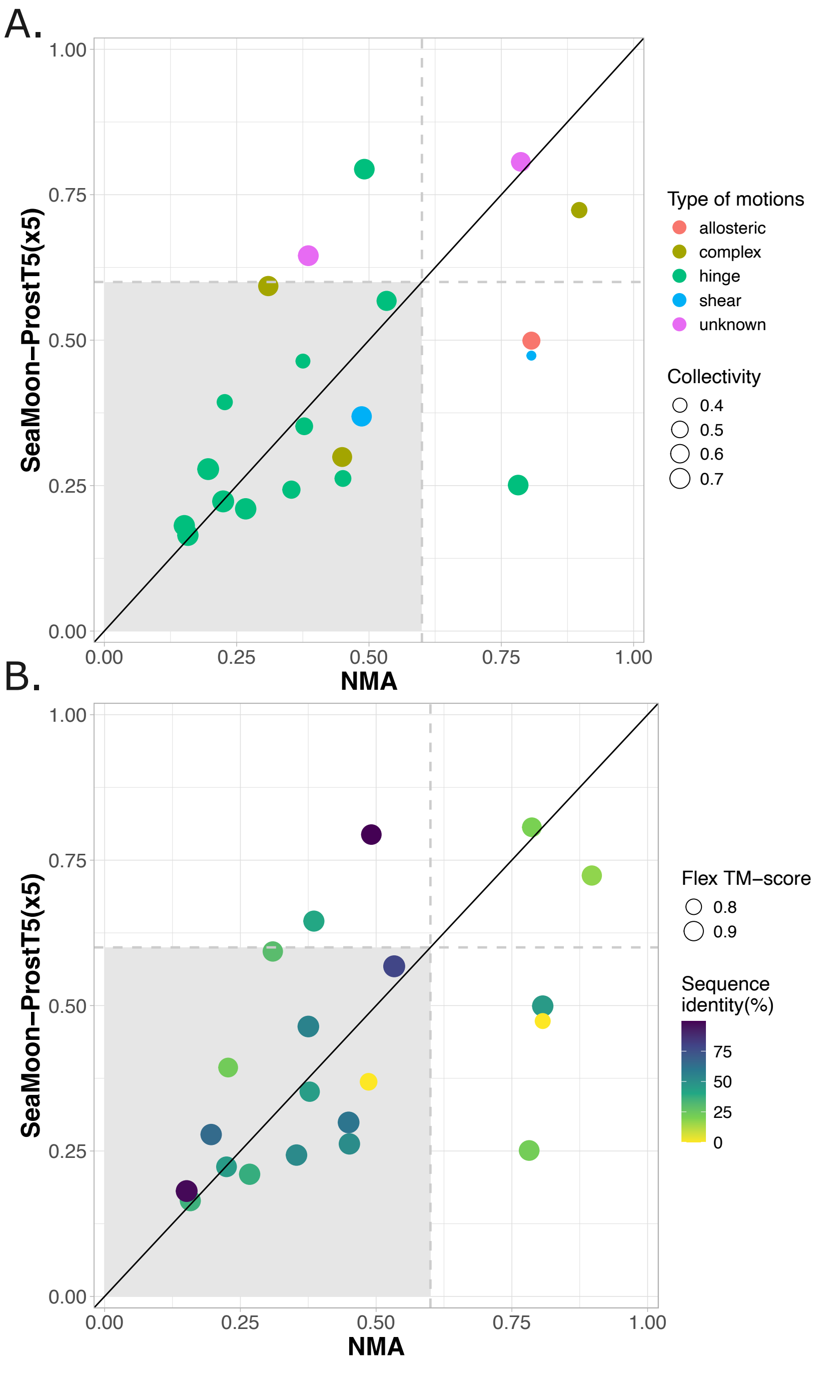


Supplemental Figure S8: **Comparison between SeaMoon-ProstT5(x5) and the NMA on the iMod benchmark, related to Figure 5.** We report the NSSE of the best-matching pair of predicted and ground-truth motions, for each method. We could assess performance on 21 proteins (see *Methods* for details on the selection): 1EX6A, 1AKEA, 8EYZG, 1LAFE, 1URPA, 1LE9A, 1EKXA, 1CKMA, 1DAPA, 1A8EA, 1MMIA, 1DMBA, 1ADBA, 1AMAA, 2J9GB, 1DPPA, 1B05A, 1AONA, 2O5CA, 1CB6A, 1OAOD, 1IG9A, 4O4BA, 1CK9A, and 1IWOA. **A.** The colour of each dot indicates the type of motion and its size is proportional to the motion collectivity. **B.** The colour gradient indicates the sequence similarity to the training set and the dot sizes are proportional to structural similarity, as measured by the Kpax flexible structural alignment algorithm.

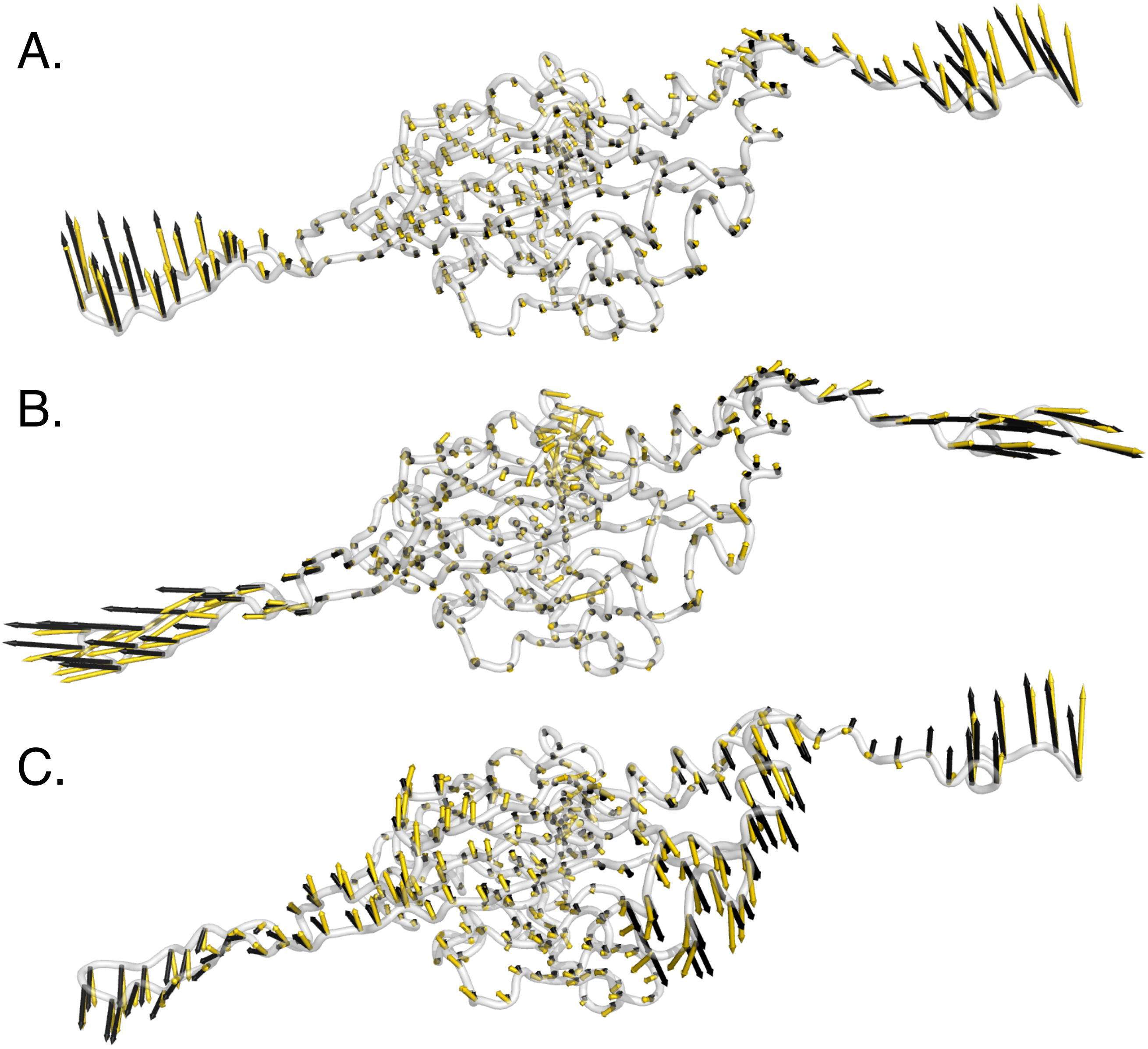


Supplemental Figure S9: **Ogopogo major capsid protein motion subspace, related to Figure 7.** Visualisation of the ground-truth (yellow) and predicted (black) motion relative directions and amplitudes for the best-matching pairs (1,1) (A), (3,2) (B) and (2,3) (C) on the reference 3D conformation, PDB code: 8ECN, chain B.

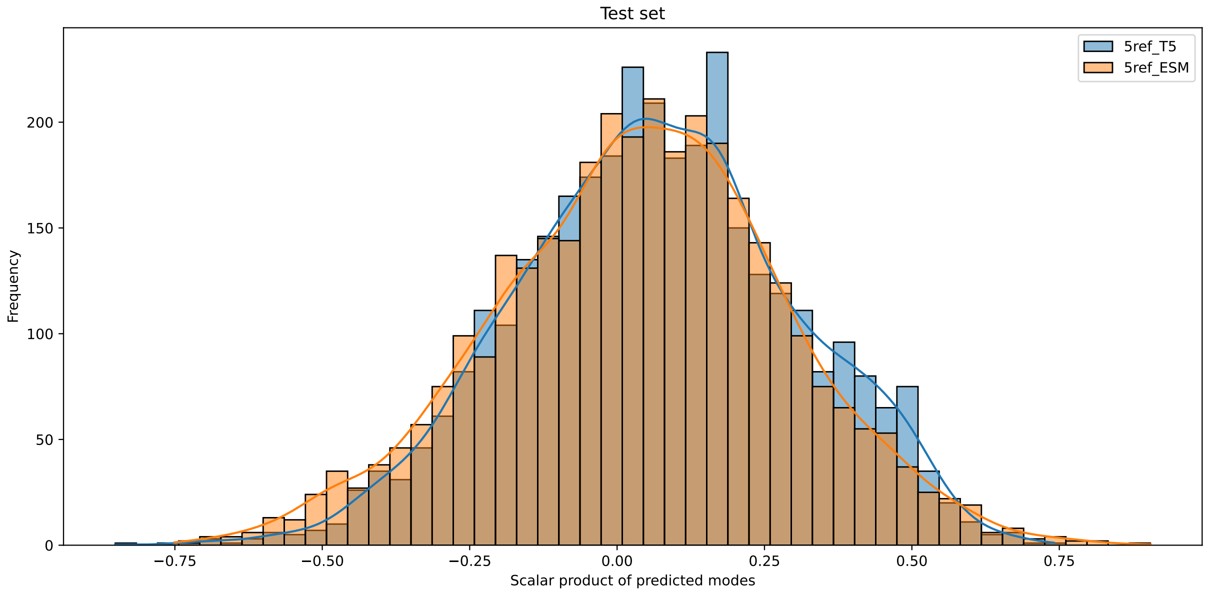


Supplemental Figure S10: **Pearson correlation computed between motions predicted by SeaMoon, related to STAR Methods.** We performed an all-to-all pairwise comparison for each protein from the test set. About 95 percent of the pairs have an absolute Pearson correlation below 0.5.
